# Comprehensive Analysis of Human Dendritic Spine Morphology and Density: Exploring Diversity of Human Dendritic Spines

**DOI:** 10.1101/2024.12.18.629169

**Authors:** Kerstin D Schünemann, Roxanne M Hattingh, Matthijs B Verhoog, Danqing Yang, Aniella Bak, Sabrina Peter, Karen M.J. van Loo, Stefan Wolking, Deborah Kronenberg-Versteeg, Yvonne Weber, Niklas Schwarz, Joseph V Raimondo, Roger Melvill, Sean Tromp, James Butler, Anke Höllig, Daniel Delev, Thomas V Wuttke, Björn M Kampa, Henner Koch

**Affiliations:** Department of Epileptology, Neurology, Uniklinik RWTH Aachen, Aachen, Germany; Neuroscience Institute, University of Cape Town, Cape Town, South Africa; Departments of Neurosurgery and of Neurology and Epileptology, Hertie-Institute for Clinical Brain Research, University of Tübingen, Tübingen, Germany; Department of Neurosurgery, RWTH Aachen University, Aachen, Germany; Systems Neurophysiology, Institute of Biology II, RWTH Aachen University, Aachen, Germany; Department of Neurosurgery, University Hospital Erlangen, Friedrich-Alexander University, Erlangen Nürnberg, Erlangen, Germany; Research Center Juelich, Institute of Neuroscience and Medicine 10, Research Center Juelich, Juelich, Germany; German Center for Neurodegenerative Diseases (DZNE) Tübingen, and Hertie Institute for Clinical Brain Research, Department of Cellular Neurology, University of Tübingen, Germany; Division of Cell Biology, Department of Human Biology, University of Cape Town, Cape Town, South Africa; Institute of Infectious Disease and Molecular Medicine, University of Cape Town, Cape Town, South Africa; Division of Neurosurgery, Department of Surgery, University of Cape Town, Cape Town, South Africa

## Abstract

Dendritic spines, small protrusions on neuronal dendrites, play a crucial role in brain function by changing shape and size in response to neural activity. So far, *in depth* analysis of dendritic spines in human brain tissue is lacking. This study presents a comprehensive analysis of human dendritic spine morphology and density using a unique dataset from human brain tissue from 27 patients (8 females, 19 males, aged 18-71) undergoing tumor or epilepsy surgery at three neurosurgery sites. We used acute slices and organotypic brain slice cultures to examine dendritic spines, classifying them into the three main morphological subtypes: Mushroom, Thin, and Stubby, via 3D reconstruction using ZEISS arivis Pro software. A deep learning model, trained on 39 diverse datasets, automated spine segmentation and 3D reconstruction, achieving a 74% F1-score and reducing processing time by over 50%. We show significant differences in spine density by sex, dendrite type, and tissue condition. Females had higher spine densities than males, and apical dendrites were denser in spines than basal ones. Acute tissue showed higher spine densities compared to cultured human brain tissue. With time in culture, Mushroom spines decreased, while Stubby and Thin spine percentages increased, particularly from 7-9 to 14 days *in vitro*, reflecting potential synaptic plasticity changes. Our study underscores the importance of using human brain tissue to understand unique synaptic properties and shows that integrating deep learning with traditional methods enables efficient large-scale analysis, revealing key insights into sex- and tissue-specific dendritic spine dynamics relevant to neurological diseases.

**New and Noteworthy:** This study presents a dataset of nearly 4,000 morphologically reconstructed human dendritic spines across different ages, gender, and tissue conditions. The dataset was further used to evaluate a deep learning algorithm for three-dimensional spine reconstruction, offering a scalable method for semi-automated spine analysis across various tissues and microscopy setups. The findings enhance understanding of human neurology, indicating potential connections between spine morphology, brain function, and the mechanisms of neurological and psychiatric diseases.

## 1 Introduction

Dendritic spines are small actin-rich protrusions along the dendrites of neurons, which contain excitatory synapses. They segregate and process synaptic inputs through AMPA and NMDA glutamate receptors, providing separate biochemical and electrical compartments for local modulation and synaptic plasticity mechanisms (Chen et al., 2011; Nimchinsky et al., 2002; Rochefort & Konnerth, 2012; Segal, 2017; Wefelmeyer et al., 2016). Spines are highly dynamic, extending and withdrawing rapidly in response to presynaptic activity (Alvarez & Sabatini, 2007; Wefelmeyer et al., 2016). The density of dendritic spines has been shown to increase during wakefulness while spines are pruned during sleep (Cirelli & Tononi, 2020; Zhou et al., 2020). Furthermore, spines have been demonstrated to increase their surface area during memory formation processes (Segal, 2017). Due to these functions, dendritic spines and their dynamics are vital for neural communication, memory formation, and learning processes.

Previous studies predominantly focused on investigating dendritic spines in neurons from animals, including rodents (Arellano et al., 2007; Koyama et al., 2015; Papa et al., 1995) and non-human primates (Chen et al., 2022; Motley et al., 2018). Studying human spines is challenging due to ethical and technical reasons but of paramount importance to improve our understanding of the human brain. Compared to rodents and primates, human pyramidal cells of the temporal cortex have stronger synaptic connections, and their spines are longer (Benavides-Piccione et al., 2002; Defelipe, 2011; Hunt et al., 2023; Iascone et al., 2020). Importantly, it has also been shown that alterations in dendritic spine density and shape are involved in neurological and psychiatric disorders such as Huntington’s disease, epilepsy, schizophrenia, bipolar disorder, and Fragile X syndrome (Fiala et al., 2002; Guidetti et al., 2001; Konopaske et al., 2014; Roberts et al., 1996). Morphological changes in dendritic spines were also identified in physiological processes such as development and aging, as spine density first increases during puberty with a subsequent constant loss of spines during adulthood (Benavides-Piccione et al., 2013; Boros et al., 2019; Petanjek et al., 2011). Notwithstanding the mounting evidence of the diversity and importance of human dendritic spines, the availability of data on this topic is still low. In addition to ethical and technical issues of gathering morphological data from human neurons, the labor-intensive analysis of the spine shape is also a limiting factor. To gain insight into the morphology of human dendritic spines, it is necessary to reconstruct numerous spines in 3D, however, manual reconstruction of dendritic spines is time-consuming and reliant on the expertise of the analyst. Algorithms can therefore aid in reconstructing larger volumes of data and annotating objects with high accuracy in a short time, e.g. by using Artificial Intelligence (AI) -generated algorithms for spine analysis (Argunsah et al., 2022; Rada et al., 2018). Reliable algorithms for spine reconstruction require extensive training datasets due to the complexity and heterogeneity of dendritic spines and limited sample sizes, numerous processing steps, or specialization in specific imaging techniques often constrain existing algorithms (Basu et al., 2018; Reberger et al., 2018; Vidaurre-Gallart et al., 2022; Vogel et al., 2023). In consequence, insights into human spine dynamics and synaptic plasticity in correlation to age, gender, and tissue condition has been scarce.

To overcome these limitations, we have gathered an extensive dataset of dendritic spines from cortical pyramidal neurons obtained from human brain surgeries. This allowed us to use machine learning-aided methods to reconstruct the shape of several thousand spines from a large cohort of human patients from several research institutions. We provide a comprehensive data set analyzing the range of spine shapes, sizes, and densities depending on dendritic region, cortical layer, gender of the human donor, and tissue condition, thereby establishing a foundation for future studies and enhancing comparability in research of human spines.

## 2 Material and Methods

### 2.1 Dataset

Brain samples from 27 patients (8 females, 19 males) aged 18 to 71 years were obtained from neurosurgical interventions at the University Hospital RWTH Aachen (Germany), University Hospital Tübingen (Germany), and the University of Cape Town (South Africa). These included tumor resections and epilepsy surgical cases (see **Table 1**). This study was performed according to the ethical and legal procedures and international regulations from the Helsinki Declaration of 1964, and its later amendments. All investigations received approval from the local ethics committees, and written consent was obtained from all patients for the scientific use of their tissue (Aachen: EK-067-20, Tübingen: 338/2016A, and Cape Town: 396/2022). Only cortical access tissue not directly affected by underlying pathologies was used. We analyzed 44 cortical layer 2/3 and 11 layer 5 pyramidal neurons. A total of 3,919 spines from these 55 neurons were examined. 11 patients (4 females, 7 males; aged 32 to 71 years) contributed 1,349 spines from 20 neurons from the University RWTH Aachen, 11 patients (2 females, 9 males; aged 18 to 67 years) contributed 1,425 spines from 14 neurons from the University of Tübingen, and 5 patients (2 females, 3 males; aged 22 to 41 years) contributed 1,145 spines from 10 neurons from the University of Cape Town. In total, 2,506 spines from acute slices (only layer 2/3) and 1413 spines from organotypic brain slice cultures (layer 2/3 and layer 5) were analyzed. See **Table 1** for detailed information on the included patients.

**Table 1.** Overview of patient specifications of samples included in dataset.

| N | Gender | Age | Pathology | Resected cortical area | DIV | Layer | Spines included |
| --- | --- | --- | --- | --- | --- | --- | --- |
| 1 | M | 22 | Epilepsy | right anterior-medial temporal | 0 | 2/3 | 354 |
| 2 | M | 26 | Tumor | left anterior temporal | 0 | 2/3 | 80 |
| 3 | F | 32 | Tumor | left temporal pole | 0 | 2/3 | 319 |
| 4 | M | 32 | Epilepsy | right temporal | 0 | 2/3 | 72 |
| 5 | F | 34 | Epilepsy | right anterior-medial temporal | 0 | 2/3 | 199 |
| 6 | M | 34 | Epilepsy | right temporal pole | 0 | 2/3 | 166 |
| 7 | F | 34 | Epilepsy | right temporal | 0 | 2/3 | 176 |
| 8 | M | 35 | Tumor | left anterior-medial temporal | 0 | 2/3 | 136 |
| 9 | F | 41 | Epilepsy | right anterior temporal | 0 | 2/3 | 304 |
| 10 | M | 47 | Epilepsy | left temporal pole | 0 | 2/3 | 46 |
| 11 | M | 48 | Epilepsy | left frontal | 0 | 2/3 | 101 |
| 12 | M | 55 | Epilepsy | right lateral temporal lobe | 0 | 2/3 | 206 |
| 13 | M | 64 | Tumor | right parietal | 0 | 2/3 | 88 |
| 14 | M | 65 | Tumor | right temporal pole | 0 | 2/3 | 74 |
| 15 | F | 67 | Tumor | right temporal pole | 0 | 2/3 | 88 |
| 16 | M | 67 | Tumor | right temporal | 0 | 2/3 | 97 |
| 17 | M | 30 | Epilepsy | right temporal | 2 | 2/3 | 91 |
| 18 | M | 18 | Epilepsy | right frontal | 3 | 2/3 | 92 |
| 19 | M | 30 | Epilepsy | left temporal | 3 | 2/3 | 47 |
| 20 | M | 30 | Tumor | left temporal | 7 | 2/3 | 184 |
| 21 | F | 40 | Epilepsy | right fronto-temporal | 8 | 2/3 | 412 |
| 22 | M | 29 | Epilepsy | left temporal | 14 | 2/3 | 67 |
| 23 | M | 47 | Epilepsy | left temporal | 3 | 5 | 49 |
| 24 | F | 32 | Tumor | left mesial temporal | 9 | 5 | 182 |
| 25 | F | 57 | Epilepsy | right temporal | 9 | 5 | 103 |
| 26 | M | 71 | Tumor | right temporo-insula | 9 | 5 | 131 |
| 27 | F | 57 | Epilepsy | right temporal | 14 | 5 | 55 |

Overview and specifics of all included patients (N). Gender is indicated as the biological sex male (M) or female (F). Age is given in years. The underlying pathology warranting surgical intervention is classified as either tumor or epilepsy origin, including malignant and benign tumors. Days *in vitro* (DIV) are given along with cortical layer localization of analyzed neurons and number of spines included in analysis.

### 2.2 Surgical Extraction

The following methods were carried out in the same way in all three centers. If there were any deviations, these are noted in each case. The tissue preparation was conducted as described previously (Bak et al., 2024; Verhoog et al., 2013). To preserve the cellular integrity of the resected tissue, care was taken during the intraoperative extraction process to minimize the use of thermal cauterization. To prevent excitotoxicity, the extracted tissue was immediately transferred into human brain slicing solution (s-aCSF), containing: 110 mM choline chloride, 26 mM NaHCO3, 1.25 mM Na2HPO4, 11.6 mM sodium ascorbate, 3.1 mM sodium pyruvate, 7 mM MgCl2, 0.5 mM CaCl2, 2.5 mM KCl, 10 mM glucose, and 1% penicillin/streptomycin/amphotericin B dissolved in Millipore water. The slicing solution was enriched with carbogen (95% O_2_, 5% CO_2_), ensuring sufficient oxygenation for the transport to the laboratories.

### 2.3 Slicing

The surgically extracted brain tissue was rinsed four times in s-aCSF and then transferred to a petri dish with ice-cold, oxygenated s-aCSF. To avoid hypoxia, the tissue was swiftly dissected into smaller blocks and trimmed to include all cortical layers and a small area of white matter. The pia mater and vessels were removed using a microdissector under a microscope. The tissue was then sectioned with a Vibratome set to 250-350 µm thickness and 0.1 mm/s cutting speed, excluding any damaged sections. To prepare the extracted tissue for further analyses, it was placed in an optimal biological environment. Firstly, the slices were transferred onto an insert using a transfer pipette. Excess fluid was aspirated from the insert. For analysis with cultured tissue, the inserts were placed on prewarmed human cerebrospinal fluid (CSF) and incubated at 37°C with 5% CO2 and 100% humidity, with a medium change every two to three days. The human CSF was collected from patients with a normal pressure hydrocephalus or idiopathic intracranial hypertension. The pooled CSF was centrifuged at 4000 rpm at 4°C for 10 minutes, then sterile-filtered through a 0,2 mm filter, and stored at -80°C.

### 2.4 Whole-Cell Patch-Clamp Recordings

The cells were filled as part of whole-cell patch clamp recordings and electrophysiological measurements. After being transferred to the recording chamber, slices were continuously perfused with carbogenated (95% O₂, 5% CO₂) recording aCSF (in mM, Aachen: 125 NaCl, 25 NaHCO_3_, 2.5 KCl, 1.25 NaH_2_PO_4_, MgCl2*6 H_2_O, 2 CaCl_2_, 25 glucose and 1% penicillin/streptomycin, Tübingen: 118 NaCl, 25 NaHCO_3_, 30 D-glucose, 1 MgCl_2_, 3 KCl, 1 NaH_2_PO_4_ and 1.5 CaCl_2_, Cape Town: 125 NaCl, 3 KCl, 1.25 NaH_2_PO4, 1MgSO4, 2 CaCl_2_, 26 NaHCO_3_ and 10 glucose) at 30-33°C. The composition of the intracellular patch solution was (in mM): K-gluconate 140, CaCl₂ 1, EGTA 10, MgCl₂ 2, Na₂ATP 4, HEPES 10. The pH was set to 7.2 with KOH/HCl, and the osmolarity was adjusted to 300 mOsm. This solution contained biocytin (5 mg/ml, Sigma, B4261). Thin-walled borosilicate glass capillaries with a resistance of 3-5 MΩ were used for the recordings. To achieve optimal filling of the entire cell, the recording was maintained for at least 20 minutes. For fixation, the slices were placed in neutral buffered 4% PFA solution at 4°C overnight after the recordings.

### 2.5 Immunohistochemistry

Slices were washed three times for ten minutes each using PBS + 0.3% Triton X-100 + 0.3% Tween-20 (PBS-T-T). Subsequently, slices were incubated in a solution made from PBS-T-T with a 1:100 ratio of 1 mg/ml Streptavidin-Cy3 (Sigma) at 4°C overnight or 1:500 AlexaFluor-488-streptavidin (Invitrogen) for 48 hours. Afterward, sections were washed again three times for 10 minutes in PBS. The Streptavidin-Cy3 (Sigma) stained slices were then incubated in 1.43 μM DAPI for a few seconds. This was followed by another washing step of three times 10 minutes in PBS. Finally, slices were mounted on microscope slides with mounting medium Fluoromount G (Invitrogen) or Vectashield (Vector Laboratories, H-1400).

### 2.6 Confocal Microscopy

Confocal microscopy was used for imaging at all three sites (Aachen and Cape Town: Zeiss LSM980 Airyscan; Tübingen: Leica TCS SP8). To ensure comparable results, imaging regions of each neuron were defined (**Figure 1A**). Neurons were manually located and focused with a 10x objective. The fluorescence was excited at 488 nm (AlexaFluor-488) and 548 nm (Cy3) using a tunable white laser, and emission was detected at 520 nm (AlexaFluor-488) and 555 nm or 615 nm (Cy3). An overview image of individual dendrites was captured. Dendrite segments of at least 40 µm were imaged to ensure a valid density analysis (**Figure 1B**). Z-stacks for spine analysis were subsequently taken with a 40x objective with a thickness of 0.22 µm to capture the entire dendrite in all dimensions. The selected segments ideally wer**e** in as few planes as possible, while maintaining imaging time and laser time at a minimum. Laser power and gain were adjusted for clear visibility of spine necks, even if slight oversaturation of the dendrite occurred. The selected dendrite was imaged with a 40x objective over a length of at least 40 µm with an electronic zoom factor of 5. Imaging settings: 1588 x 1588 pixels, 16-bit depth, Speed 6, Average 0, unidirectional. Post-processing was performed using the ‘Joint Deconvolution’ method with approximately 20 iterations to capture all recording details.

**Figure 1.**
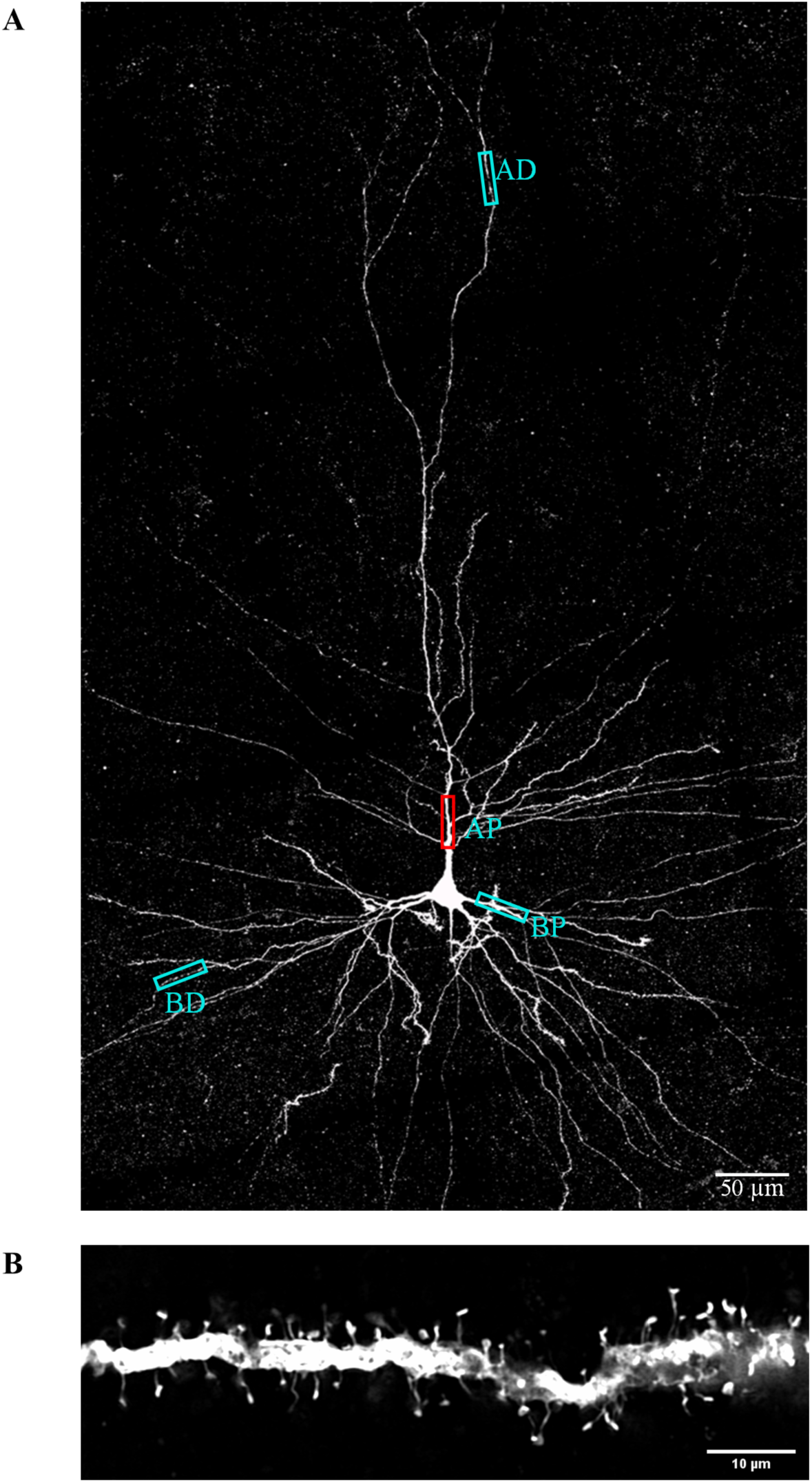
Confocal Imaging Regions of Pyramidal Neurons. (A) Exemplary maximum intensity projection confocal image of a human layer 2/3 pyramidal cell of the temporal cortex. Regions of interest (cyan boxes) were defined to facilitate further analysis: apical distal (AD) as a terminal segment of the apical tuft; apical proximal (AP) approximately 70 µm from the soma; basal proximal (BP) as second or third order segments; basal distal (BD) as terminal segments of basal dendrite. (B) Example of a single optical plane of a confocal image of the apical proximal (AP) dendrite (red box) from the neuron shown in (A).

### 2.7 3D-Reconstruction

To analyze the morphology of dendritic spines, 3D reconstructions of confocal images were generated using ZEISS arivis Pro (Version 4.1.1, Carl Zeiss Microscopy, 2023). The Magic Wand Tool with a 1% tolerance was used to manually detect and reconstruct the dendritic spines. Pixels of the spines that were not detected correctly by the semi-automatic tool were corrected manually. A merged object combining the dendrite and spines was visualized in 3D mode. Using the cutting tool in 3D mode, spines were separated from the dendrite for accurate analysis of spine necks. The software calculated the surface area and volume for each 3D object. In 2D mode, manual measurements of total length (nm), diameter of the spine head (nm), and diameter of the spine neck (nm) were performed using anatomical structures as reference points. The diameter of the head was determined at the widest point, and the diameter of the neck was determined approximately at the halfway point, avoiding the transition from the head to the neck to obtain accurate measurements. Spine density per μm was calculated based on the number of all identifiable spines on at least a 40 µm dendritic stretch.

### 2.8 Deep Learning Algorithm

To facilitate the evaluation of our large data set, we trained a deep learning model using the Zeiss arivis Cloud (Carl Zeiss Microscopy, 2023), a cloud-based AI platform. First, confocal images were uploaded in .tiff or .czi format to the cloud storage. An annotation tool was used to define training categories for the algorithm: dendrite, spines, and background. Annotations were made in a single plane. Extensive annotations were added to the images to instruct the algorithm on the specific elements it should or should not identify in the following reconstructions. Special attention was given to areas with low fluorescent signals, particularly spine necks. The training process was carried out on the arivis Cloud platform, which operated within an Azure Kubernetes Service (AKS) cluster, and trained on 39 datasets (13 Cape Town, 13 Tübingen, 13 Aachen) to handle heterogeneous data sources. This setup facilitated efficient deployment, management, and scaling of containerized applications. The platform automated the entire machine learning process, encompassing data preparation, model training, deployment, and management. Continuous model improvement was achieved through retraining based on previous iterations, ensuring reliable and efficient segmentation results. The system utilized Azure Kubernetes Service, microservices architecture, event-driven communication, and Azure Machine Learning for dendritic spine segmentation. The trained model was imported into arivis Vision4D and implemented as a reconstruction workflow.

Its performance was compared to 380 manually reconstructed spines from 12 patients (4 females, 8 males), blinded to age. Filtering thresholds of 1 µm² and 30 µm² surface area were applied to remove irrelevant objects. False positives, representing background noise or dendrite segments misclassified as spines, constituted 49.6% of the detected objects. The model also identified extra spines missed during manual reconstruction. True-positive spines were manually measured for length, head, and neck diameter, while the model automatically calculated volume and surface area. Both manual and model-based reconstructions used the same imaging datasets for comparison. The high rate of false positive spines was reduced quickly to 0 by manually deciding whether it was a true spine or not and excluding false positives. In the direct comparison, not all spines found by the manual reconstruction were detected by the model. The rate of undetected spines was 3.6%. The rate of extra spines, referring to objects found by the model and not marked in manual reconstruction, but which were real spines, was 6.2%. Due to the lower fluorescent signal of spine necks, the algorithm did not always identify the entire spine with head and neck. In total, 79.1% of actual spines were completely depicted with head and neck, while a spine head without a connecting neck was found in every case. In some cases, objects were reconstructed by the model that combined two or more spines in one object due to their close spatial proximity. The rate of these overlapping objects was 12%. The model performed with a sensitivity of 61%, recall of 95%, and an F1-score of 74%. The model-based reconstruction significantly reduced the time required for manual reconstruction, decreasing the time per spine from 90 seconds to 35 seconds.

Following the examination of the quantity of spines detected by the deep learning tool, their quality was assessed. A total of 380 spines (patients n = 12) were examined in direct comparison, and only spines detected in both the manual and the model-based reconstruction were included. Extra spines and false positives were excluded from the analysis. The parameters surface area, volume, length, diameter head, and diameter neck (**Figure 2A**) were examined. No significant differences were observed in the parameters surface area (**Figure 2B**) (manual: 7.06 ± 0.34 µm², model: 7.32 ± 0.41 µm², p > 0.99), volume (**Figure 2C**) (manual: 0.63 ± 0.05 µm³, model: 0.77 ± 0.06 µm³, p = 0.32) and length (**Figure 2D**) (manual: 1301 ± 44 nm, model: 1182 ± 40 nm, p = 0.91). There was also no statistically significant difference for diameter head (**Figure 2E**) (manual: 511 ± 12 nm, model: 526 ± 15 nm, p = 0.12) and diameter neck (**Figure 2F**) (manual: 258 ± 10 nm, model: 271 ± 10 nm, p = 0.84).

**Figure 2.**
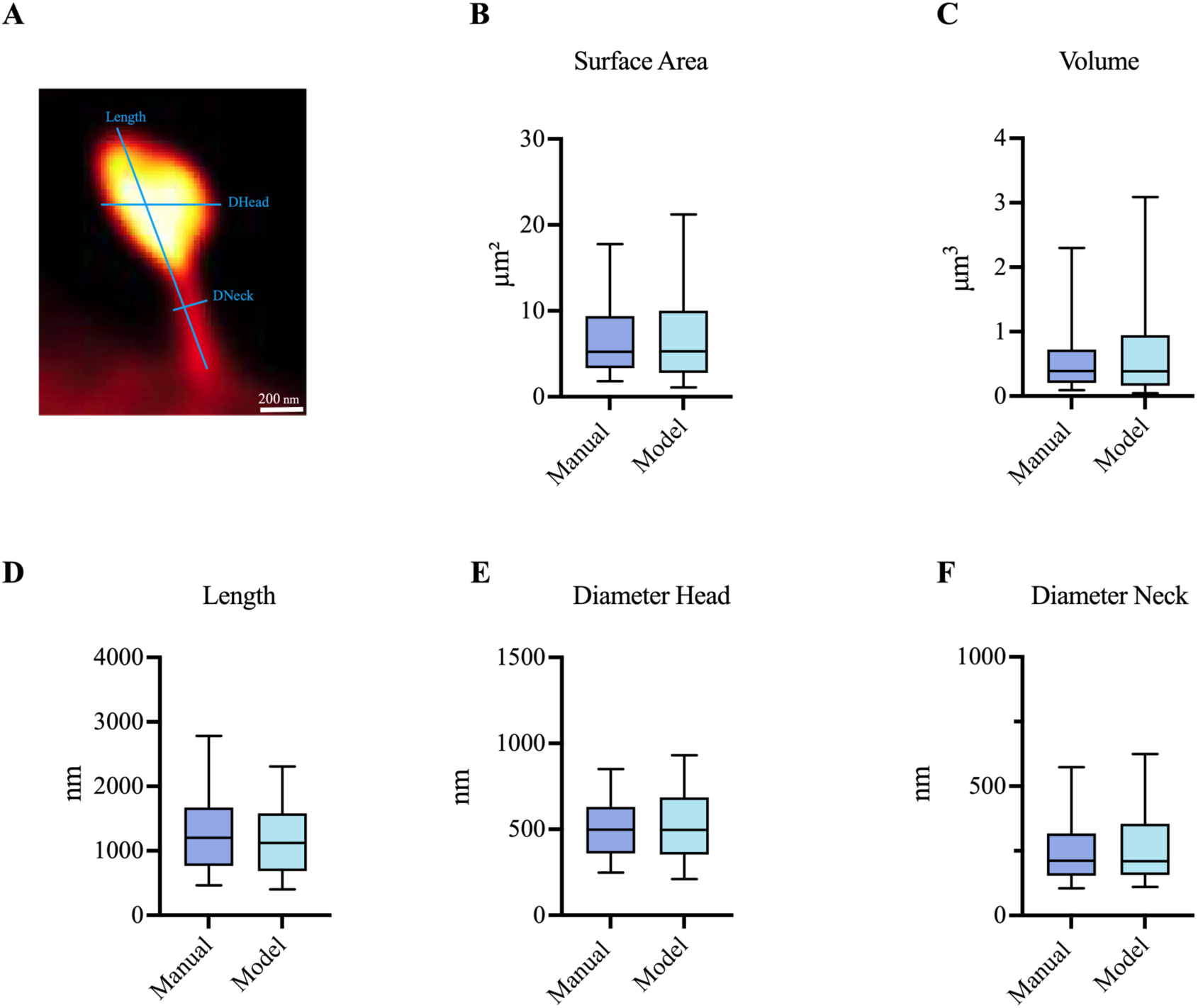
Efficiency evaluation of deep learning model. (A) Depiction of manual measurements of spines in one plane of the confocal image. Performance of the deep learning tool in surface area (B), volume (C), length (D), diameter head (E), and diameter neck (F) as compared to the manual reconstruction. Significance tested utilizing Friedman test with p < 0.05. Line in the middle of the box shows the mean, whiskers indicate the 5;95% confidence interval.

### 2.9 Spine Classes

Spines were classified into four morphological subtypes, i.e. Mushroom, Thin, Filopodia, and Stubby using established criteria with ratios adapted from (Harris et al., 1992) (**Figure 3A**). The Diameter Neck/Diameter Head ratio (**Figure 3B**) and Diameter Head/Spine Length ratio were determined (**Figure 3C**), and the cut-off value was set at 0.8 for both. Mushroom spines had larger spine heads than necks, with a cut-off value at < 0.8 Diameter Neck/Diameter Head Ratio (**Figure 3H**). Thin spines had narrow heads and long, slender necks. Spines were classified as ‘Thin’ if their length exceeded the diameter of the head (Diameter Head/Spine Length < 0.8) and the Diameter Neck/Diameter Head ratio was > 0.8 (**Figure 3I**). Filopodia spines are thin spines with a total spine length > 3 μm (Qiao et al., 2016). Stubby spines had no or very weakly pronounced necks. Spines were categorized as ‘Stubby’ if Diameter Neck/Diameter Head Ratio was > 0.8 and (Diameter Neck/Length Ratio > 0.8 OR Spine Length < 700nm) (**Figure 3J**).

**Figure 3.**
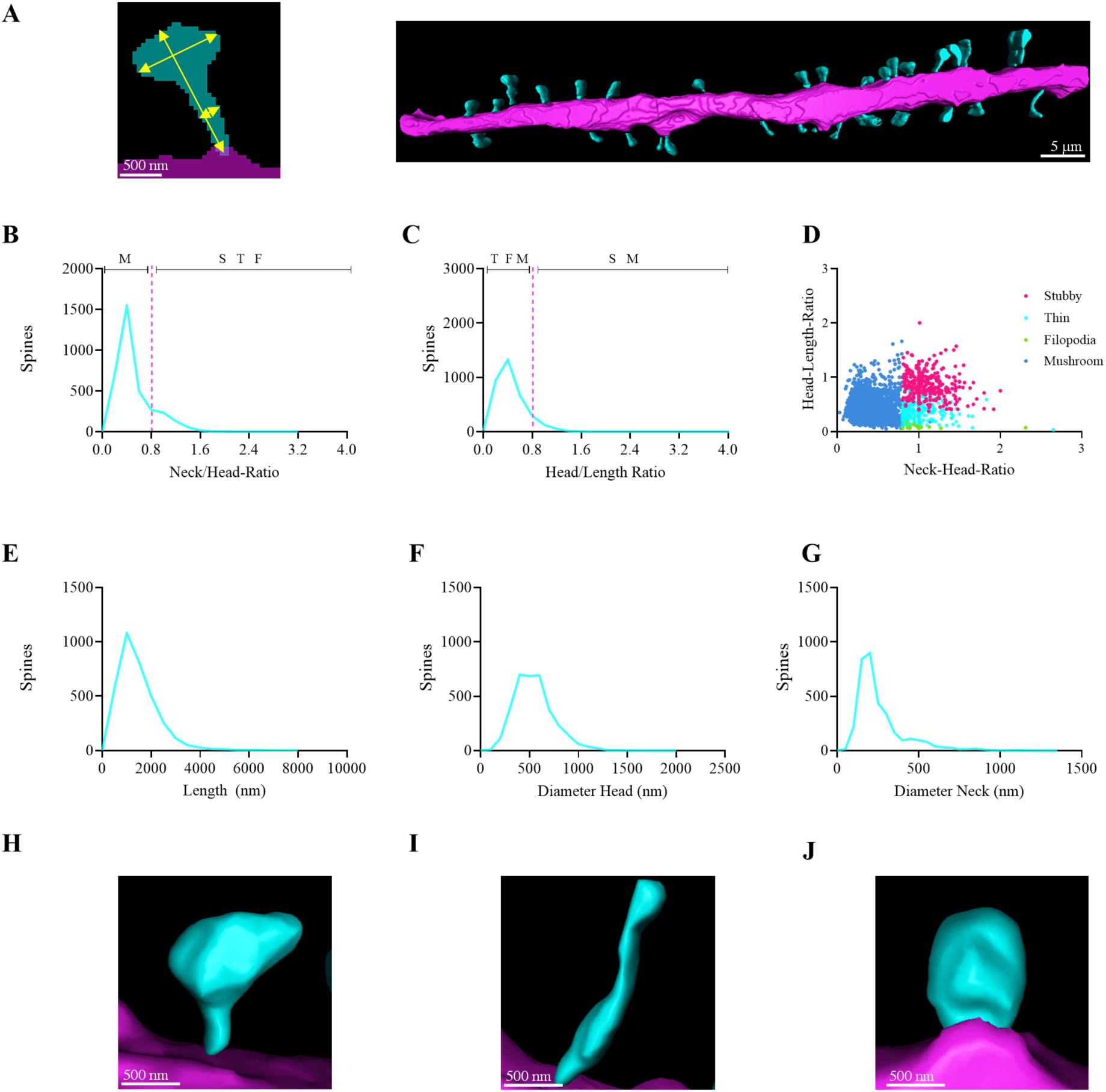
Spine Parameter Distribution. (A) Depiction of manual 2D measurement of spine length, head diameter, and neck alongside a manual 3D reconstruction of an exemplary apical dendrite of a 32-year-old male patient. The dendrite is shown in pink, the spines are shown in cyan. Diameter Neck/Diameter Head ratio and Diameter Head/Spine Length ratio were determined. (B) Distribution of Neck/Head ratio, cut-off value at 0.8. (C) Distribution of Head/Length ratio, cut-off value at 0.8. (D) Neck/Head ratio and Head/length ratio plotted against each other. Spine classes visualized in different colors, namely Mushroom („M“, blue), Thin („T“, cyan), Stubby („S“, magenta), and Filopodia („F“, green). (E) Distribution of Spine Length. (F) Distribution of head diameter. (G) Distribution of neck diameter. Exemplary reconstructions shown for (H) Mushroom, (I) Thin, and (J) Stubby subclasses.

### 2.10 Statistical Analysis

Statistical analysis was conducted using GraphPad Prism (Version 10.1, GraphPad Software, Boston, Massachusetts, USA). Because of non-normally distributed data, Kruskal-Wallis, Mann-Whitney test, or two-tailed Friedman test with Dunn’s multiple comparison test were performed. The null hypothesis stated that there was no significant difference (α = 0.05) between manual reconstruction and reconstruction by the algorithm. The percentages of undetected spines, false-positive spines, extra spines, overlapping spines, and complete spines were evaluated for each version. Morphological parameters, including surface area, volume, length, diameter head, and diameter neck, were examined.

## 3 Results

### 3.1. Spine Density in Adult Human Pyramidal Cells

To gain an initial insight into the differences in human spines, we first analyzed their density. The analysis of 56 dendrites from layer 2/3 neurons originating from acute brain slices from females (n= 5, 32 to 67 years, mean 41.6 years) versus 68 dendrites from males (n= 11, 22 to 67 years, mean 45 years) indicated a significant difference with a higher spine density in females (female: 0.51 ± 0.03 spines/μm, male: 0.46 ± 0.05 spines/μm, p = 0.03) (**Figure 4A**). A significant difference in dendritic density was observed between apical (47 dendrites, n (females)= 23, n (males)= 24) and basal dendrites (25 dendrites, n (females)= 10, n (males)= 15) (**Figure 4B**). The density of dendritic spines was significantly higher in the apical dendrites compared to the basal areas (apical: 0.68 ± 0.08 spines/μm, basal: 0.40 ± 0.04 spines/μm, p = 0.01). There was no difference between layer 2/3 (21 dendrites) and layer 5 dendrites (18 dendrites) (layer 2/3: 0.41 ± 0.04 spines/μm, layer 5: 0.36 ± 0.05 spines/μm, p = 0.48) (**Figure 4C**). To see if there is an influence of age on spine density, we formed 3 age groups: 20-40 years, 41-50 years and 50-67 years. Due to the shown difference in dendritic spine density between men and women, each age group was first categorized by gender. No significant difference was found for females (group 20-40 years (n = 27) 0.54 ± 0.05 spines/μm, group 41-50 years (n = 7) 0.69 ± 0.07 spines/μm, group 51-67 years (n = 2) 0.83 ± 0.08 spines/μm, p (20-40 vs. 41-50 years) 0.17, p (41-50 vs. 51-67 years) > 0.99) and males (group 20-40 years (n = 17) 0.66 ± 0.14 spines/μm, group 41-50 years (n = 9) 0.36 ± 0.08 spines/μm, group 51-67 years (n = 19) 0.48 ± 0.08 spines/μm, p (20-40 vs. 41-50 years) > 0.99, p (41-50 vs. 51-67 years) > 0.99) . Therefore, both genders were combined to represent a larger number of patients. Again, no significant difference in spine density was found **(Figure 4D)** (group 20-40 years (n = 41) 0.57 ± 0.07 spines/μm, group 41-50 years (n = 16) 0.50 ± 0.07 spines/μm, group 51-67 years (n = 21) 0.51 ± 0.08 spines/μm, p (20-40 vs. 41-50 years) > 0.99, p (41-50 vs. 51-67 years) > 0.99). A significantly higher density was also found for dendrites from acute tissue (78 dendrites, n (females) = 33, n (males) = 45) compared to cultured tissue (50 dendrites, n (females)= 23, n (males) = 27) (acute tissue: 0.55 ± 0.04 spines/μm, cultured tissue: 0.38 ± 0.02 spines/μm, p < 0.0001) (**Figure 4E**). In summary, a higher dendritic spine density was found in women as compared to men, in apical dendrites as compared to basal dendrites, and in dendrites from acute tissue as compared to cultured tissue.

**Figure 4.**
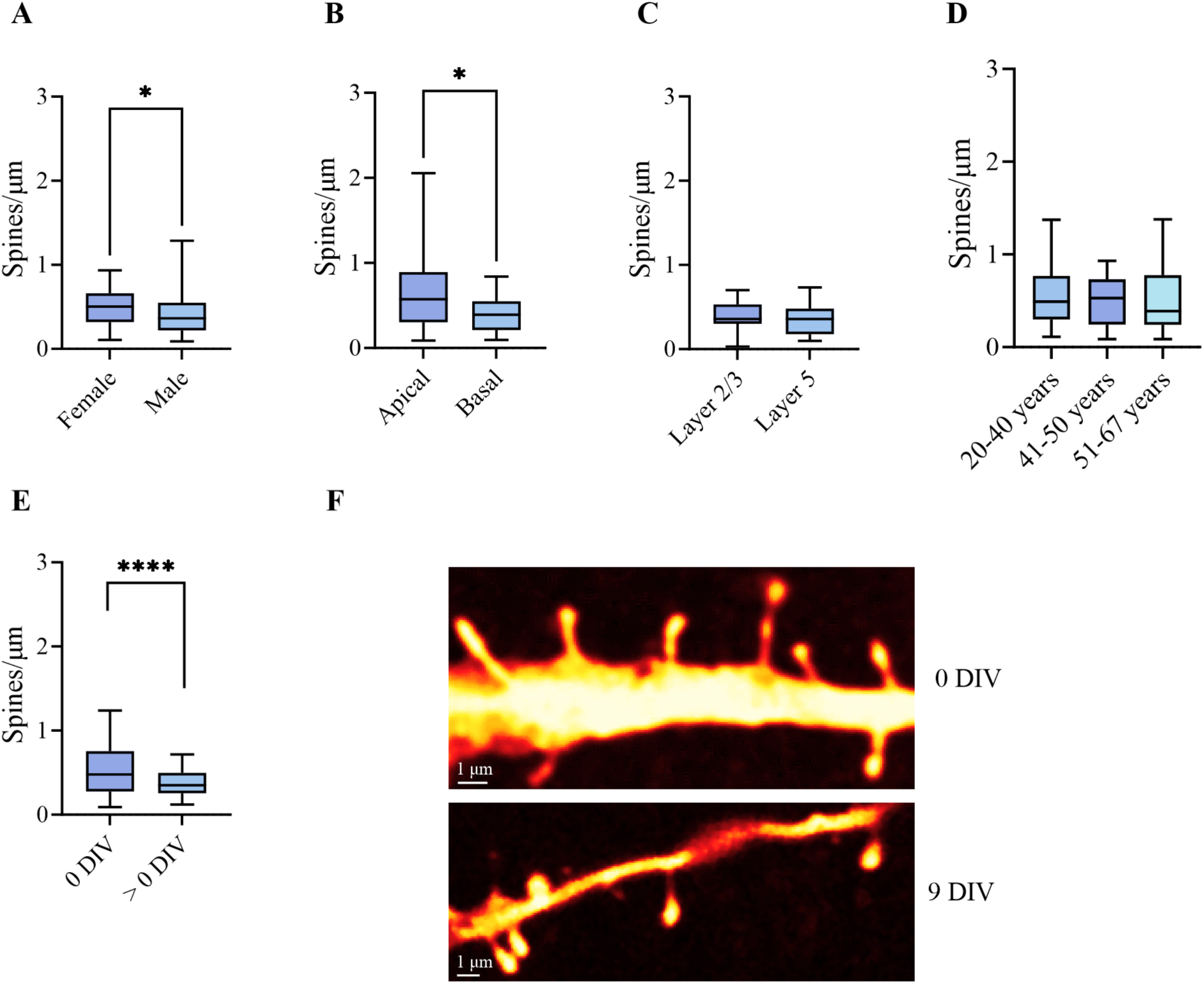
Comparison of spine densities. (A) Comparison of spine density between males and females, (B) apical and basal dendrites, (C) Layer 2/3 and Layer 5, (D) different age groups and (E) acute (0 DIV) cultured (> 0 DIV) tissue. (F) Exemplary image of an apical dendrite originating from acute tissue (0 DIV) and an image of an apical dendrite originating from cultured tissue (9 DIV). * = p ≤ 0.05, ** = p ≤ 0.01 and **** = p < 0.0001 for Kruskal-Wallis test. Line in the middle of the box shows the mean, whiskers indicate the 5;95% confidence interval.

### 3.2. Spine Class Distribution in Adult Human Pyramidal Cells

Following the spine classification according to their morphological parameters in 3D reconstruction, the subsequent step was to analyze the distribution of the spine classes as a percentage of the total number of spines. z-Test was used to test significance. Only 1 Filopodia spine could be detected in layer 2/3 pyramidal cells from acute tissue. Therefore, this class was not considered further for the distribution. First, we analyzed whether differences were present between tissue from tumor (T) and epilepsy surgery (E). No significant difference was observed in the distribution (Mushroom 84% (T) vs. 85% (E), Stubby 11% (T) vs. 10% (E), Thin 5% (T) vs. 5% (E)). Therefore, the spines were pooled for the following subgroup analysis. The distribution of spine types in layer 2/3 pyramidal cells of female patients was found to be 83% Mushroom, 12% Stubby, and 5% Thin spines. In male subjects, the distribution was 86% Mushroom, 10% Stubby, and 4% Thin spines. A z-test revealed no significant differences in the distribution (**Figure 5A**). No significant difference in the distribution of spine classes between apical (a) and basal (b) dendrites could be shown (Mushroom 86% (a) vs. 81% (b), Stubby 9% (a) vs. 15% (b), Thin 5% (a) vs. 4% (b)) (**Figure 5B**). Similarly, there was no difference in the distribution between layer 2/3 and layer 5 spines (Mushroom 75% (2/3) vs. 80% (5), Stubby 8% (2/3) vs. 4% (5), Thin 17% (2/3) vs. 16% (5)) (**Figure 5C**). There were no differences in the spine class distribution between the different age groups 20-40 years (Mushroom: 87%, Stubby 9%, Thin 4%), 41-50 years (Mushroom: 82%, Stubby 11%, Thin 7%) and 51-67 years (Mushroom: 85%, Stubby 9%, Thin 6%) **(Figure 5D).** It can be stated that, Mushroom spines constitute the majority of dendritic spines. The data indicate that Thin spines are more prevalent than Stubby spines in layers 2/3 and 5. It is important to note that this analysis was conducted using spines derived from cultured tissue, whereas the comparison of apical/basal and female/male was analyzed from acute tissue.

**Figure 5.**
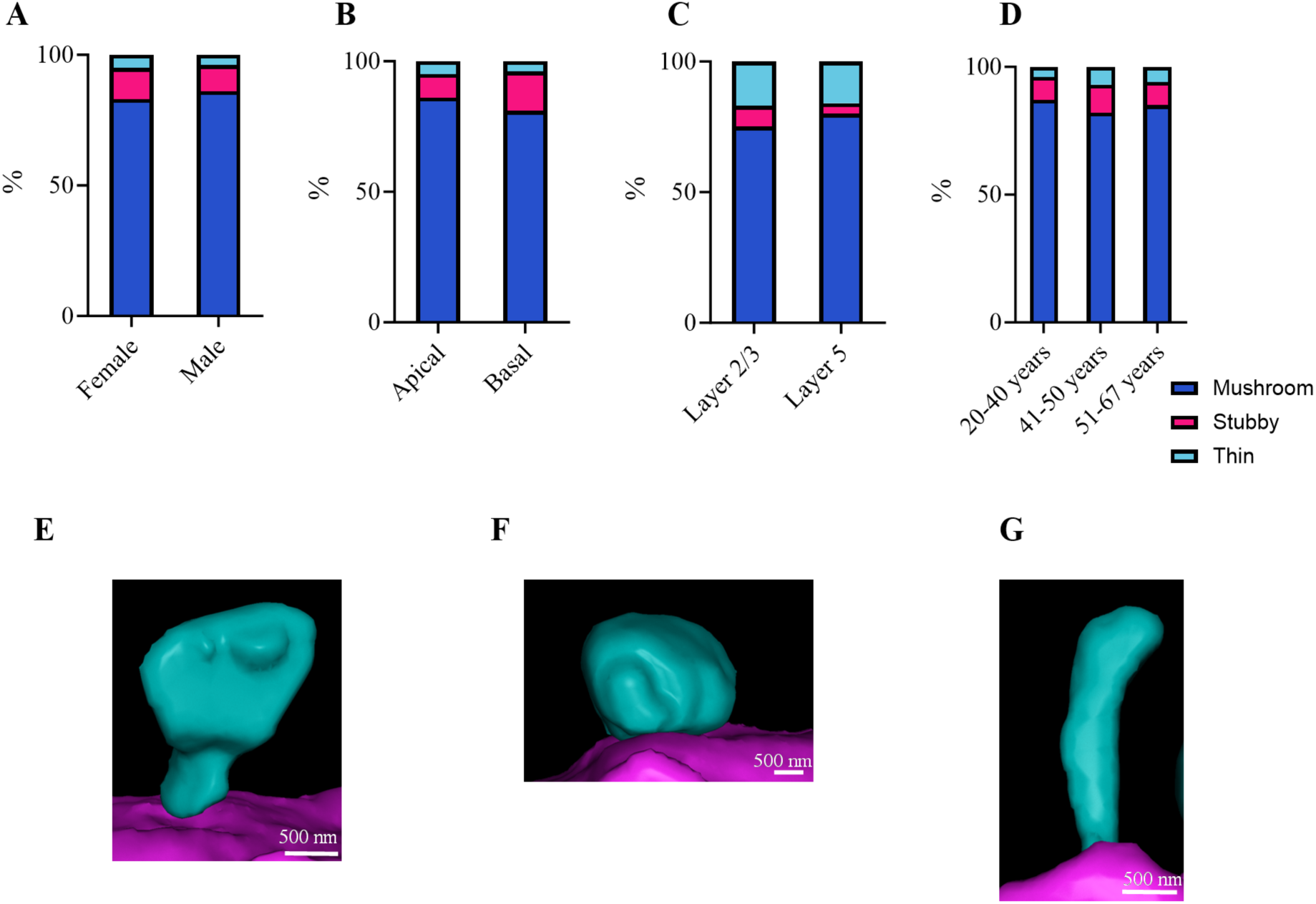
Comparison of Spine Subclass Proportions. Stacked bar charts illustrate the distribution of the three different spine classes: Mushroom (blue), Stubby (magenta), and Thin (cyan). Distribution is given for (A) Female vs. Male, (B) Apical vs. Basal, (C) Layer 2/3 vs. Layer 5, and (D) three different age groups. Exemplary 3D-reconstructions of a (E) Mushroom spine, (F) Stubby spine, and (G) Thin spine.

### 3.3. Differences in Morphological Parameters of Spine Subclasses

To obtain a more detailed analysis of the spine morphology we also analyzed whether there are morphological differences between the three spine classes as compared between the subgroups. For a better overview, the exact values for all classes can be found for Mushroom spines in **Table 2**, Stubby spines in **Table 3** and Thin Spines in **Table 4**. Mushroom spines obtained from tissue samples during tumor surgeries (n = 660) and epilepsy surgeries (n = 1150) were subjected to detailed analysis. No statistically significant differences were observed across any of the examined parameters, including surface area, volume, length, and head and neck diameter. Consequently, the spines were pooled for subsequent analysis. Similarly, Stubby spines from access tissue obtained from tumor (n = 88) or epilepsy surgeries (n = 144) were analyzed. No statistically significant difference was detected for any parameter (p > 0.99). Thin spines from access tissue obtained from tumor operations (n = 36) and epilepsy operations (n = 63) were initially analyzed. The results demonstrated that there was no statistically significant difference for all parameters. There was no significant difference for any of the subgroups, except for neck diameter, which is larger for layer 2/3 than layer 5 thin spines.

**Table 2.** Comparison of Mushroom Spine Parameter for different subgroups.

| **<u>Mushroom (Spines)</u>** | **<u>Surface ($\mu\text{m}^2$)</u>** | **<u>Volume ($\mu\text{m}^3$)</u>** | **<u>Length (nm)</u>** | **<u>DiaHead (nm)</u>** | **<u>DiaNeck (nm)</u>** |
| --- | --- | --- | --- | --- | --- |
| Female (n= 900) | 5. 87 ± 0.12 | 0.48 ± 0.01 | 1412 ± 23 | 535 ± 6 | 190 ± 2 |
| Male (n= 910) | 6.76 ± 0.15 | 0.60 ± 0.02 | 1500 ± 23 | 554 ± 6 | 197 ± 3 |
| p-value | > 0.99 | >0.99 | > 0.99 | > 0.99 | > 0.99 |
| Apical (n= 1267) | 6.21 ± 0.11 | 0.51 ± 0.01 | 1490 ± 20 | 544 ± 5 | 189 ± 2 |
| Basal (n= 398) | 5.21 ± 0.15 | 0.4 ± 0.02 | 1314 ± 30 | 509 ± 9 | 185 ± 3 |
| p-value | > 0.99 | > 0.99 | > 0.99 | > 0.99 | > 0.99 |
| Layer 2/3 (n= 473) | 8.99 ± 0.25 | 0.84 ± 0.03 | 1649 ± 35 | 578 ± 8 | 264 ± 4 |
| Layer 5 (n= 284) | 8.86 ± 0.30 | 0.80 ± 0.04 | 2065 ± 63 | 619 ± 15 | 232 ± 6 |
| p-value | > 0.99 | > 0.99 | > 0.99 | > 0.99 | > 0.99 |
| 20-40 years (n= 1165) | 5.69 ± 0.12 | 0.48 ± 0.02 | 1422 ± 20 | 537 ± 6 | 199 ± 2 |
| 41-50 years (n= 288) | 5.74 ± 0.02 | 0.54 ± 0.02 | 1422 ± 38 | 577 ± 11 | 206 ± 4 |
| p-value | > 0.99 | > 0.99 | > 0.99 | > 0.99 | > 0.99 |
| 51-67 years (n= 388) | 8.38 ± 0.22 | 0.69 ± 0.02 | 1517 ± 37 | 545 ± 9 | 184 ± 3 |
| p-value | > 0.99 | > 0.99 | > 0.99 | > 0.99 | > 0.99 |

Comparison of spine parameters surface, volume, length, head diameter, and neck diameter between subgroups female vs. male, apical vs. basal, and layer 2/3 vs. layer 5 for Mushroom spines. Statistical significance tested via Kruskal-Wallis test followed by Dunn's multiple comparison test. Results are shown as mean ± standard error of the mean.

**Table 3.** Comparison of Stubby Spine Parameter for different subgroups.

| **<u>Stubby (Spines)</u>** |  |  |  |  |  |
| --- | --- | --- | --- | --- | --- |
| Female (n= 128) | 3.23 ± 0.18 | 0.25 ± 0.02 | 575 ± 13 | 573 ± 14 | 482 ± 13 |
| Male (n= 104) | 3.93 ± 0.03 | 0.36 ± 0.04 | 597 ± 17 | 496 ± 19 | 522 ± 18 |
| p-value | > 0.99 | > 0.99 | > 0.99 | > 0.99 | > 0.99 |
| Apical (n= 139) | 3.48 ± 0.18 | 0.28 ± 0.02 | 575 ± 13 | 478 ± 14 | 501 ± 13 |
| Basal (n= 72) | 3.15 ± 0.19 | 0.24 ± 0.02 | 591 ± 19 | 478 ± 20 | 471 ± 19 |
| p-value | > 0.99 | > 0.99 | > 0.99 | > 0.99 | > 0.99 |
| Layer 2/3 (n= 51) | 5.7 ± 0.45 | 0.48 ± 0.06 | 609 ± 23 | 595 ± 23 | 611 ± 27 |
| Layer 5 (n= 19) | 5.64 ± 0.55 | 0.50 ± 0.06 | 593 ± 35 | 608 ± 39 | 666 ± 54 |
| p-value | > 0.99 | > 0.99 | > 0.99 | > 0.99 | > 0.99 |
| 20-40 years (n= 121) | 3.38 ± 0.27 | 0.29 ± 0.04 | 603 ± 15 | 476 ± 16 | 507 ± 16 |
| 41-50 years (n= 39) | 2.98 ± 0.23 | 0.22 ± 0.03 | 543 ± 29 | 455 ± 22 | 519 ± 23 |
| p-value | > 0.99 | > 0.99 | > 0.99 | > 0.99 | > 0.99 |
| 51-67 years (n= 41) | 4.76 ± 0.39 | 0.42 ± 0.05 | 561 ± 25 | 493 ± 29 | 550 ± 24 |
| p-value | > 0.99 | > 0.99 | > 0.99 | > 0.99 | > 0.99 |

Comparison of spine parameters surface, volume, length, head diameter, and neck diameter between subgroups female vs. male, apical vs. basal, and layer 2/3 vs. layer 5 for Stubby spines. Statistical significance tested via Kruskal-Wallis test followed by Dunn's multiple comparison test. Results are shown as mean ± standard error of the mean.

**Table 4.**
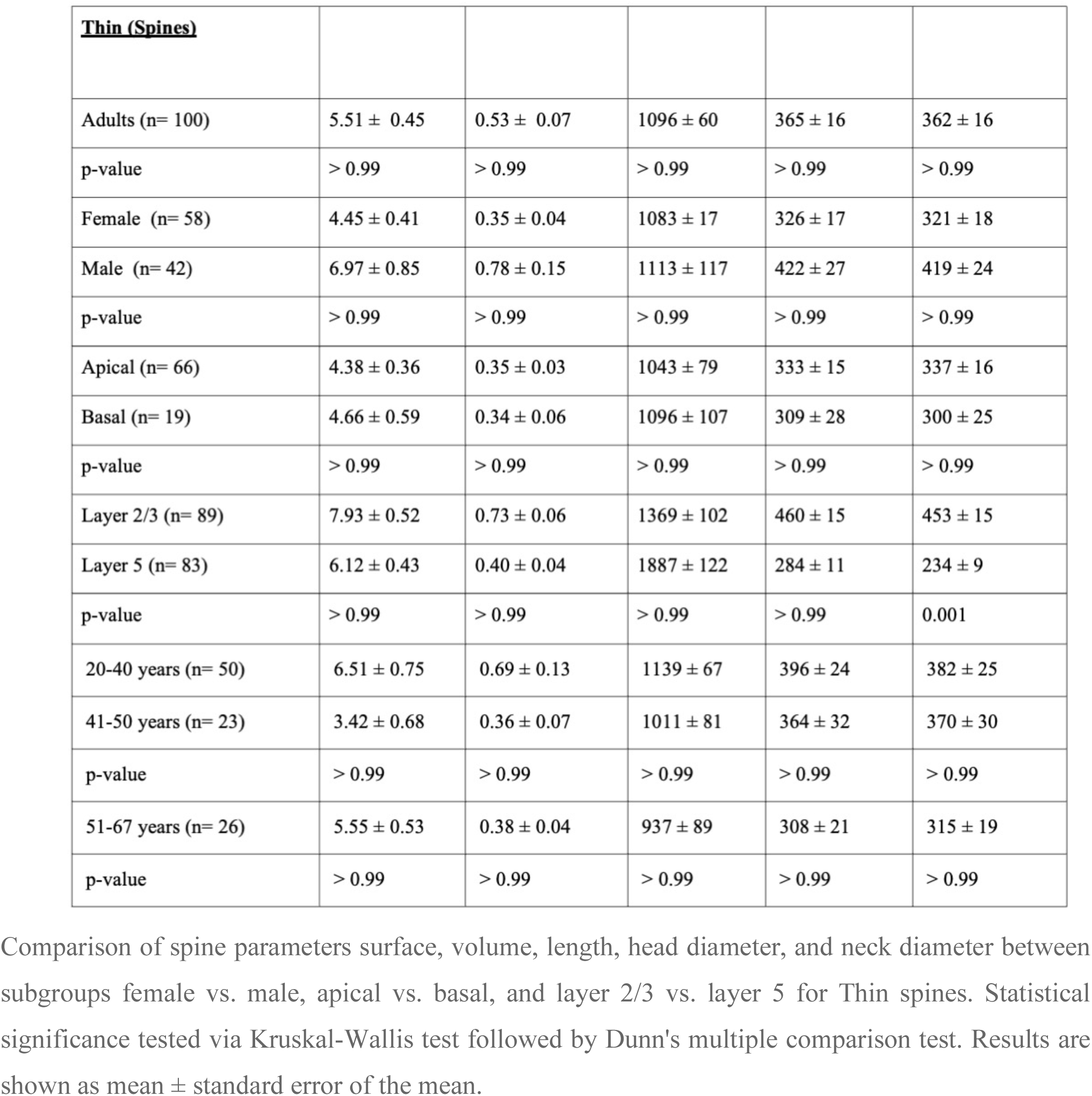
Comparison of Thin Spine Parameter for different subgroups.

| **<u>Thin (Spines)</u>** |  |  |  |  |  |
| --- | --- | --- | --- | --- | --- |
| Adults (n= 100) | 5.51 ± 0.45 | 0.53 ± 0.07 | 1096 ± 60 | 365 ± 16 | 362 ± 16 |
| p-value | > 0.99 | > 0.99 | > 0.99 | > 0.99 | > 0.99 |
| Female (n= 58) | 4.45 ± 0.41 | 0.35 ± 0.04 | 1083 ± 17 | 326 ± 17 | 321 ± 18 |
| Male (n= 42) | 6.97 ± 0.85 | 0.78 ± 0.15 | 1113 ± 117 | 422 ± 27 | 419 ± 24 |
| p-value | > 0.99 | > 0.99 | > 0.99 | > 0.99 | > 0.99 |
| Apical (n= 66) | 4.38 ± 0.36 | 0.35 ± 0.03 | 1043 ± 79 | 333 ± 15 | 337 ± 16 |
| Basal (n= 19) | 4.66 ± 0.59 | 0.34 ± 0.06 | 1096 ± 107 | 309 ± 28 | 300 ± 25 |
| p-value | > 0.99 | > 0.99 | > 0.99 | > 0.99 | > 0.99 |
| Layer 2/3 (n= 89) | 7.93 ± 0.52 | 0.73 ± 0.06 | 1369 ± 102 | 460 ± 15 | 453 ± 15 |
| Layer 5 (n= 83) | 6.12 ± 0.43 | 0.40 ± 0.04 | 1887 ± 122 | 284 ± 11 | 234 ± 9 |
| p-value | > 0.99 | > 0.99 | > 0.99 | > 0.99 | 0.001 |
| 20-40 years (n= 50) | 6.51 ± 0.75 | 0.69 ± 0.13 | 1139 ± 67 | 396 ± 24 | 382 ± 25 |
| 41-50 years (n= 23) | 3.42 ± 0.68 | 0.36 ± 0.07 | 1011 ± 81 | 364 ± 32 | 370 ± 30 |
| p-value | > 0.99 | > 0.99 | > 0.99 | > 0.99 | > 0.99 |
| 51-67 years (n= 26) | 5.55 ± 0.53 | 0.38 ± 0.04 | 937 ± 89 | 308 ± 21 | 315 ± 19 |
| p-value | > 0.99 | > 0.99 | > 0.99 | > 0.99 | > 0.99 |

Comparison of spine parameters surface, volume, length, head diameter, and neck diameter between subgroups female vs. male, apical vs. basal, and layer 2/3 vs. layer 5 for Thin spines. Statistical significance tested via Kruskal-Wallis test followed by Dunn's multiple comparison test. Results are shown as mean ± standard error of the mean.

### 3.4. Spines from acute vs. cultured slices in adult human pyramidal cells

To obtain reliable insights into the morphology of human dendritic spines and possible changes occurring during the culturing process, we analyzed whether spines obtained from acute slices differed from spines from cultured brain slices. Four periods were defined for this purpose: 0 DIV, 2-3 DIV, 7-9 DIV and 14 DIV. A total of 893 spines from cultured brain tissue from 6 patients (n(male) = 5, n(female)= 1) were included in the analysis. There were 223 spines 2-3 DIV from 3 patients (all male), 602 spines 7-9 DIV (n (male)= 1, n(female)= 1), and 68 spines 14 DIV (n(male)= 1). To see whether the subgroups could be pooled in further analyses, the percentage distribution of the spine classes was first evaluated (**Figure 6A**). Mushroom spines from acute tissue accounted for 86%, Stubby spines for 9%, and Thin spines for 9%. In comparison, Mushroom spines accounted for 87%, Stubby spines for 9%, and Thin spines for 4% in dendrites that were 2-3 DIV. Using a z-test, no significant difference in the distribution could be shown. For spines from 7-9 DIV tissue, a proportion of 75% Mushroom spines, 11% Stubby spines, and 13% Thin spines could be identified. In comparison with the percentage distribution of spines from 2-3 DIV, there was therefore a significantly lower proportion of Mushroom spines in 7-9 DIV neurons and a significantly higher proportion of Thin spines compared to neurons that were 2-3 DIV. For spines from 14 DIV, 60% Mushroom spines, 21% Stubby spines, and 19% Thin spines could be analyzed. Compared to 7-9 DIV, the proportion of Mushroom spines was significantly lower, but the proportion of Stubby and Thin spines was not significantly different. However, when the proportions of spine classes from 14 DIV and 0 DIV were compared, a clear picture emerges: the percentage of Mushroom spines significantly decreases over time in culture while the percentage of Stubby and Thin spines significantly increases. Likewise, Stubby spines are represented in acute tissue with a higher proportion than thin spines. In cultured tissue, this difference is almost equalized.

**Figure 6.**
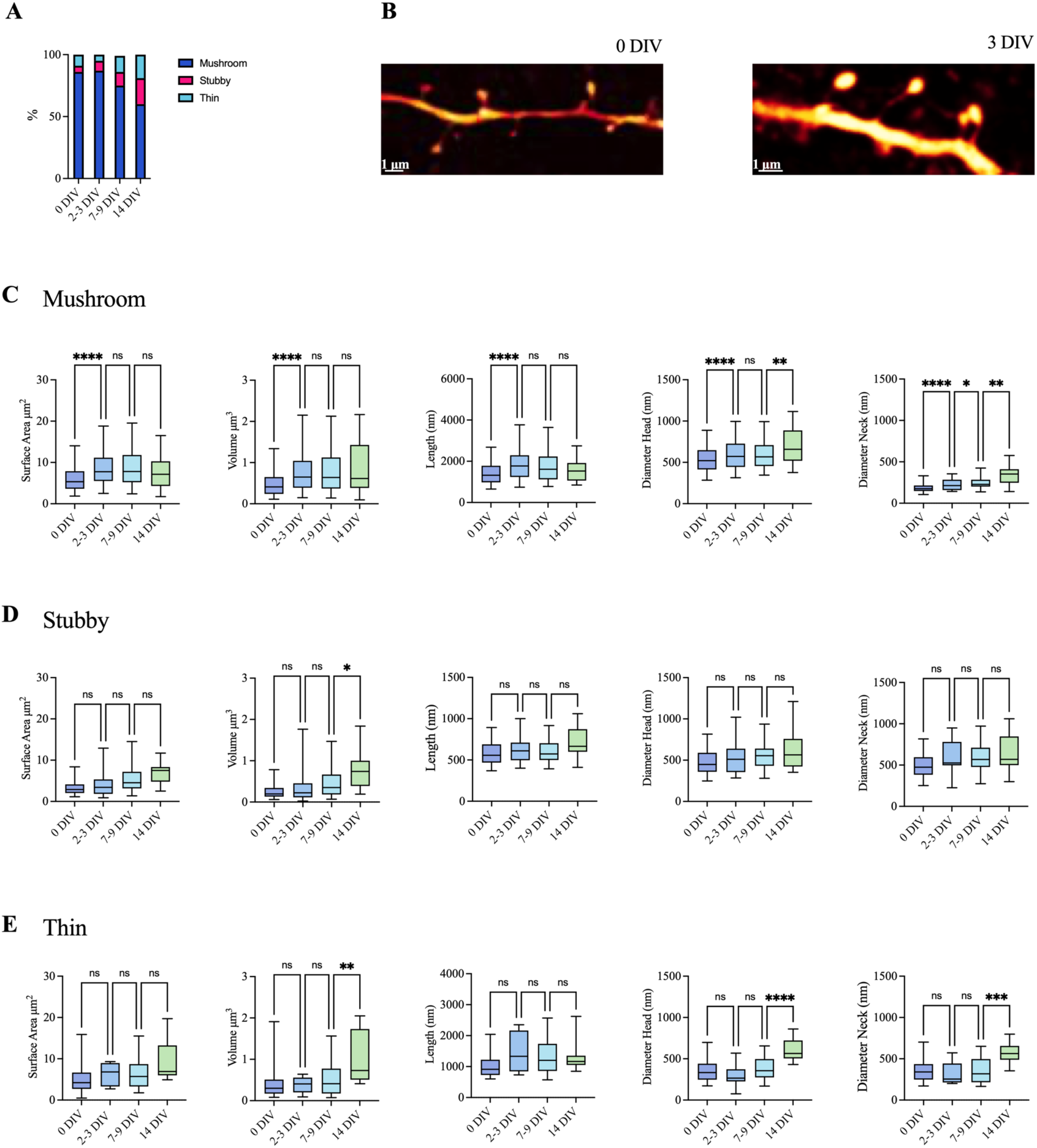
(A) Spine class distribution for periods acute, 2-3 DIV, 7-9 DIV, and 14 DIV. (B) Exemplary confocal microscopy image of a dendrite in acute tissue (left) and in tissue 3 days *in vitro* (right). Parameters surface area, volume, length, head diameter, and neck diameter were analyzed for (C) Mushroom spines, (D) Stubby spines, and (E) Thin spines. * = p ≤ 0.05, ** = p ≤ 0.01 and **** = p < 0.0001 for Kruskal-Wallis test. Line in the middle of the box shows the mean, whiskers indicate the 5;95% confidence interval.

Since a difference in the percentage distribution of the three different spine classes had already been demonstrated, this classification was also maintained for the morphological analysis. First, the Mushroom spines were examined, including 1841 spines from acute tissue, 194 from 2-3 DIV, 707 from 7-9 DIV, and 56 from 14 DIV (**Figure 6C**). For a better overview, the exact values for all classes can be found in **Table 5**. It was shown that all parameters for spines from 2-3 DIV were significantly higher than those from acute tissue. No differences in morphological parameters were observed between Mushroom spines from 2-3 DIV and 7-9 DIV. However, Mushroom spines from 14 DIV showed a larger head and neck diameter compared to those from 7-9 DIV. In summary, Mushroom spines at 2-3 DIV exhibit the greatest morphological difference from acute tissue, characterized by an overall increase in size and shape.

**Table 5.** Comparison of Spine Class Parameters between Acute and Cultured Tissue.

| **<u>Mushroom (Spines)</u>** | **<u>Surface ($\mu\text{m}^2$)</u>** | **<u>Volume ($\mu\text{m}^3$)</u>** | **<u>Length (nm)</u>** | **<u>DiaHead (nm)</u>** | **<u>DiaNeck (nm)</u>** |
| --- | --- | --- | --- | --- | --- |
| Acute (n= 1841) | 6.31 ± 0.10 | 0.54 ± 0.01 | 1456 ± 16 | 545 ± 4 | 194 ± 2 |
| 2-3 DIV (n= 194) | 9.02 ± 0.2 | 0.81 ± 0.04 | 1906 ± 70 | 610 ± 14 | 231 ± 6 |
| 7-9 DIV (n= 707) | 9.02 ± 0.20 | 0.83 ± 0.03 | 1800 ± 34 | 598 ± 8 | 249 ± 4 |
| 14 DIV (n= 56) | 7.7 ± 0.62 | 0.89 ± 0.1 | 1598 ± 90 | 700 ± 29 | 338 ± 17 |
| p-value 0 vs. 2-3 DIV | < 0.0001 | < 0.0001 | < 0.0001 | < 0.0001 | < 0.0001 |
| p-value 0 vs. 7-9 DIV | < 0.0001 | < 0.0001 | < 0.0001 | < 0.0001 | < 0.0001 |
| p-value 0 vs. 14 DIV | 0.76 | 0.0001 | 0.37 | < 0.0001 | < 0.0001 |
| p-value 2-3 vs. 7-9 DIV | > 0.99 | > 0.99 | 0.63 | > 0.99 | 0.35 |
| p-value 7-9 vs. 14 DIV | 0.39 | > 0.99 | > 0.99 | 0.006 | 0.002 |
| **<u>Stubby (Spines)</u>** |  |  |  |  |  |
| Acute (n= 232) | 3.54 ± 0.17 | 0.30 ± 0.02 | 584 ± 11 | 483 ± 11 | 500 ± 11 |
| 2-3 DIV (n= 19) | 4.22 ± 0.70 | 0.377 ± 0.09 | 623 ± 32 | 534 ± 42 | 600 ± 48 |
| 7-9 DIV (n= 92) | 5.66 ± 0.40 | 0.49 ± 0.05 | 611 ± 16 | 562 ± 19 | 598 ± 22 |
| 14 DIV (n= 19) | 7.17 ± 0.60 | 0.77 ± 0.1 | 716 ± 43 | 625 ± 52 | 657 ± 48 |
| p-value 0 vs. 2-3 DIV | > 0.99 | > 0.99 | > 0.99 | > 0.99 | 0.18 |
| p-value 0 vs. 7-9 DIV | < 0.0001 | < 0.0001 | 0.73 | 0.0008 | 0.0003 |
| p-value 0 vs. 14 DIV | < 0.0001 | < 0.0001 | 0.01 | 0.03 | 0.0008 |
| p-value 2-3 vs. 7-9 DIV | 0.255 | 0.77 | > 0.99 | > 0.99 | > 0.99 |
| p-value 7-9 vs. 14 DIV | 0.115 | 0.01 | 0.17 | > 0.99 | > 0.99 |
| **<u>Thin (Spines)</u>** |  |  |  |  |  |
| Acute (n= 99) | 5.53 ± 0.45 | 0.53 ± 0.07 | 1056 ± 46 | 366 ± 16 | 362 ± 16 |
| 2-3 DIV (n= 10) | 6.32 ± 0.79 | 0.38 ± 0.06 | 1455 ± 193 | 296 ± 42 | 314 ± 43 |
| 7-9 DIV (n= 146) | 6.51 ± 0.34 | 0.54 ± 0.04 | 1326 ± 50 | 383 ± 13 | 361 ± 14 |
| 14 DIV (n= 14) | 9.29 ± 1.22 | 0.99 ± 0.16 | 1277 ± 113 | 610 ± 35 | 361 ± 14 |
| p-value 0 vs. 2-3 DIV | 0.94 | > 0.99 | 0.166 | > 0.99 | 576 ± 34 |
| p-value 0 vs. 7-9 DIV | 0.09 | > 0.99 | 0.0006 | > 0.99 | > 0.99 |
| p-value 0 vs. 14 DIV | 0.003 | 0.0007 | 0.17 | < 0.0001 | > 0.99 |
| p-value 2-3 vs. 7-9 DIV | > 0.99 | 0.07 | > 0.99 | 0.54 | > 0.99 |
| p-value 7-9 vs. 14 DIV | 0.09 | 0.005 | > 0.99 | < 0.0001 | 0.0001 |

Comparison of spine parameters surface, volume, length, head diameter, and neck diameter between subgroups acute, 2-3 DIV, 7-9 DIV, and 14 DIV for each spine type. Statistical significance tested via Kruskal-Wallis test followed by Dunn's multiple comparison test. Results are shown as mean ± standard error of the mean.

Stubby spines were also examined (**Figure 6D**). For this purpose, 232 Stubby spines from acute tissue, 19 from 2-3 DIV, 92 from 7-9 DIV, and 19 from 14 DIV were analyzed. The subgroups are quite small in absolute terms, as the proportion of Stubby spines is only around 15% of all spines. Over time from 0 to 14 DIV, a significant increase could only be shown for the volume from 7-9 DIV to 14 DIV. All other parameters were not significantly different. However, it could be shown that the parameters differed significantly from 0 DIV compared with 7-9 DIV except for the length. Here, the parameters surface, volume, diameter head, and neck were significantly larger than in stubby spines from acute tissue. When comparing 0 DIV with 14 DIV, a significant increase in all parameters, including length, was also shown. Overall, it can therefore be said that all parameters increase over time, that they do not differ significantly in direct comparison day by day but do in comparison to 0 DIV tissue.

For the class of Thin spines (**Figure 6E)**, 99 spines from acute tissue, 10 spines from 2-3 DIV, 146 spines from 7-9 DIV, and 14 spines from 14 DIV could be included. For the period 0 to 2-3 DIV and 2-3 DIV to 7-9 DIV, no significant difference was observed for any of the parameters analyzed. For the period 7-9 DIV to 14 DIV, a significantly larger volume, diameter head, and neck could be shown for the thin spines from 14 DIV. There was no significant difference between 0 DIV and 7-9 DIV, but a significant difference in Surface, Volume, and Diameter Head could be shown in comparison with 14 DIV. In summary, morphological changes of Thin spines tend to take place from the second week *in vitro*.

## 4 Discussion and Conclusion

This study provides a comprehensive analysis of dendritic spine morphology and density. Our unique dataset comprised brain tissue samples from 27 patients who underwent neurosurgery for tumors or epilepsy, collected from three hospitals: University Hospital RWTH Aachen, University Hospital Tübingen, and the University of Cape Town. By combining data from acute and cultured tissue, this study offers an *in-depth* examination of spine morphology across diverse biological contexts, patient demographics, and experimental conditions.

The use of human brain tissue is critical for understanding the human brain, as findings from animal models and cell culture systems often do not translate directly to human biology. Despite significant investments in preclinical studies using animal models (Seyhan, 2019), the failure to translate these findings into effective human treatments is well-documented (Dhir et al., 2020; Franke et al., 1996; Macdonald et al., 2011; Wahlgren et al., 1999). This highlights the urgent need for studies examining the human brain’s functional and morphological properties at the cellular and circuit levels (Lee et al., 2024). Although the fundamental motifs of synaptic connectivity between pyramidal cells and interneurons in the human brain share many features with those of other species, such as comparable cortical GABAergic interneurons, excitatory pyramidal cells, and similar synaptic connectivity patterns (Molnar et al., 2016), distinct properties of human synapses and their connections necessitate direct human studies (Peng et al., 2024; Yang et al., 2024). Recent studies investigating human brain slices have started to explore the unique properties of human neuronal dendrites, which are structurally and functionally distinct from those in rodents (Eyal et al., 2016; Gidon et al., 2020; Mohan et al., 2015). The increased size and functional differences of these dendrites may cause significant alterations in signal propagation from synapses to the soma and back due to voltage attenuation along these extensive, cable-like structures (Beaulieu-Laroche et al., 2018; Gooch et al., 2022; Stuart & Sakmann, 1994). These biophysical differences could impact synaptic integration and plasticity differently in humans compared to animal models, underscoring the importance of detailed analysis of dendritic spine morphology to gain a better understanding of the human brain.

Our study reveals significant differences in dendritic spine density based on sex, dendrite type, and tissue condition, providing new insights into the structural diversity of human cortical neurons. Female patients exhibited a higher spine density than males, aligning with findings from animal studies suggesting hormonal influences. In animal studies, spine density decreases with lower estrogen and testosterone levels while high estrogen levels shift spine morphology to smaller spines, indicating a close relationship between sex hormones and spine morphology (Brandt et al., 2020; Chen et al., 2013; Garelick & Swann, 2014; Jabra et al., 2024; Shors et al., 2001; Woolley et al., 1990; Young et al., 2014). These hormonal effects may also be relevant in humans and could account for the higher dendritic spine density observed in females. Future studies incorporating estrogen data would allow a more precise analysis of gender differences. Such an approach would require ethical considerations, including additional blood draws, as well as accounting for fluctuations in estrogen levels over the menstrual cycle. While logistically challenging, this line of research could yield critical insights into gender-specific neurological health. Moreover, although men generally have larger cerebrums, the gray matter is denser in females (Filipek et al., 1994; Goldstein et al., 2001; Passe et al., 1997; Salminen et al., 2022). This denser gray matter could contribute to the greater spine density observed in women, potentially influencing synaptic connectivity and neuronal circuitry. The implications of these structural differences extend to various neurological and psychiatric conditions, where differences in synaptic density and distribution could affect disease susceptibility, progression, and treatment response.

Our analysis found no significant sex differences in the distribution of spine types (Mushroom, Stubby, Thin). This lack of difference suggests that while overall spine density differs by sex, the relative distribution of spine types remains consistent. An essential characteristic of dendritic spines is their ability to dynamically regulate both volume and density. Stimuli that trigger long-term potentiation and depression result in alterations in spine volume through the activation of NMDA receptors (Matsuzaki et al., 2004; Tazerart et al., 2020; Zhou et al., 2004), which in turn may affect spine volume by facilitating the recruitment of AMPA receptors. In the central nervous system, immature synapses on thin filopida spines exhibit a low AMPA to NMDA receptor ratio, which increases during development and learning (Malinow & Malenka, 2002; Vardalaki et al., 2022). Whereas, mushroom spines have been shown to be indicators of stronger synaptic connections due to their larger head size and more significant postsynaptic density (Bourne & Harris, 2008; Nusser et al., 1998). Therefore, the distribution of spine morphologies in the individual brain may also reflect the extent of previously learned skills and memories. It has also been well described in humans and animal models, that spine morphology and density is altered in neurological diseases such as schizophrenia and autism (Penzes et al., 2011). While spines are lost in Alzheimer’s disease, the spine density is usually increased in autism spectrum disorder. Together, these findings indicate that other factors, including lifestyle, genetic background, neurological pathologies or non-hormonal physiological differences, may influence spine morphology.

Interestingly, our findings indicate that dendritic spine density and morphology remain consistent throughout adulthood in our human dataset, aligning with previous observations in limited human studies. These studies suggest a decrease in spine density from young to adult ages, with density stabilizing after approximately age 40 (Benavides-Piccione et al., 2013; Jacobs et al., 1997), which is in line with our results. Spine head volume has also been shown to remain constant with age, which is consistent with our results (Lenz et al., 2024). In addition, we were able to show that the parameters length, surface and diameter of the neck also remain constant with increasing age. Animal studies have documented a notable decrease in spine density from youth to adulthood (Duan et al., 2003; Kabaso et al., 2009; Page et al., 2002; Peters et al., 1998). Notably, these animal studies primarily compared young and aged groups, whereas our analysis focuses on adult human tissue only. To gain a more comprehensive understanding, future studies involving samples from a broader age range, including infant and pediatric patients, are needed. This could provide valuable insights into the neurobiological factors influencing dendritic spine stability and their implications for cognitive function and neurological disease susceptibility throughout the human lifespan.

We also observed differences in dendritic spine density between the apical and basal parts of the neuron, consistent with previous studies showing more clustered spines in apical dendrites (Anton-Sanchez et al., 2017; Benavides-Piccione et al., 2013). Our analysis demonstrated that these more complex patterns were not due to differences in spine morphology, which did not show any significant differences. Also, we found no differences in spines on pyramidal neurons of layer 2/3 compared to layer 5. However, these neurons consist of two major classes including intratelencephalic (IT-type) neurons in layer 2/3 and 5, which mediate cortico-cortical communication, and layer 5 pyramidal tract (PT-type) neurons that project outside the telencephalon to the brainstem and spinal cord (Harris & Shepherd, 2015). And these different cell types express distinct roles during decision-making tasks in mice (Musall et al., 2023). While it has been shown that spine density changes on IT-type neurons in frontal cortex during adolescence (Delevich et al., 2020), these experiments depend on projection target specific labeling of these neuron types which is currently not possible in humans. Staining neurons during patch-clamp recordings, however, allows retrieval of the transcriptomics of the neuron which would allow identifying the cell type (Berg et al., 2021).

To address the need for efficient and reliable morphological analysis, we evaluated an AI-based method for automatic detection and 3D reconstruction of dendritic spines across data from the three neurosurgery hospital sites. The deep learning model effectively detected dendritic spines, with false positives easily curated manually. Spine shapes, including spine heads, were faithfully reconstructed. Manual intervention was only necessary to add spine necks, which are challenging to detect due to low fluorescence, but this still reduced the time required for analysis. The model accurately determined morphological parameters such as head diameter, surface area, and volume, similar to manually reconstructed spines. Our 3D-based approach distinguishes itself from existing methods relying on 2D data or specific microscopy adaptations, which have limitations in handling 3D data (Levet et al., 2020; Reberger et al., 2018; Smirnov et al., 2018; Vogel et al., 2023; Xiao et al., 2018). While our algorithm achieved similar accuracy to these methods, challenges related to image quality, especially along the Z-axis, remain. Future efforts should focus on enhancing algorithm stability and integrating diverse methodologies to fully understand the morphology-function relationship of dendritic spines.

In our analysis, we classified dendritic spines into the three established categories: mushroom, stubby, and thin spines, following established methods (Harris et al., 1992). We set quantitative criteria for distinguishing these classes to eliminate potential biases from purely visual categorization. Unlike other studies e.g. (Young et al., 2014), which use absolute cut-off values, we adopted a ratio-based approach to enhance comparability not only within human samples but also across species. Several software packages for automatic spine classification are already available, which consider key parameters beyond individual spine dimensions for categorizing spines (Ghani et al., 2017; Gilles et al., 2024; Shi et al., 2014; Swanger et al., 2011). However, we propose that rigid classification schemes may not fully capture physiological conditions, as spine morphology likely represents a continuum rather than discrete categories. This idea is supported by previous studies in rodents, showing that spines exist along a morphological spectrum, observed in both live and fixed tissue (Arellano et al., 2007; Berry & Nedivi, 2017; Loewenstein et al., 2015; Tonnesen et al., 2014; Yuste & Bonhoeffer, 2004). We are available to provide our dataset on request for a specific comparison. Additionally, it is important to recognize the limitations imposed by confocal microscopy on spine morphology analysis due to resolution constraints. Spines are typically no longer than 4000 nm, with spine heads often measuring below 1000 nm. Confocal microscopes provide a resolution of approximately 200–300 nm, potentially obscuring smaller or thinner spines, as well as the spine neck region, which is crucial for many classification schemes. Future studies may benefit from AI-assisted classification that considers a continuum of parameters rather than rigid categories or clusters. Incorporating synaptic activity in addition to morphological features could provide a more holistic approach, allowing for spine categorization based on both structure and function. This approach may offer valuable new insights into the relationships between appearance and functionality of dendritic spines.

Using human organotypic slice cultures to measure function and morphology has been shown to be a valid tool in several recent studies (Lee et al., 2023; Schwarz et al., 2019; Ting et al., 2018). Nevertheless, concerns exist that cell properties may change significantly during the culturing process. However, most electrophysiological and morphological features, such as firing rate, action potential patterns, and network activity, seem to remain stable over time (Lee et al., 2023; Schwarz et al., 2019; Ting et al., 2018). A slight but significant increase in the resting potential of pyramidal cells has been reported particularly between days two and three in culture (Schwarz et al., 2019; Ting et al., 2018). To address whether changes occur in spines during culture, we included an analysis of cells recorded from acute and cultured slices over 14 days *in vitro*. The temporal analysis of spines from cultured brain slices revealed significant changes in spine morphology over time. Specifically, mushroom spines decreased, while stubby and thin spines increased significantly from 7 to 14 DIV, reflecting potential synaptic plasticity changes. It is well-documented that thin spines can mature into mushroom spines as animals or humans engage in new learning experiences (Bourne & Harris, 2007; Hayashi & Majewska, 2005). Given that learning processes cannot occur in culture, it may be reasonable to hypothesize that the dynamics are reversed, leading to a reduction in mushroom spines. These findings underscore the importance of considering culture duration in experimental designs when studying spine morphology. In addition to time spent *in vitro*, the potential effects of anesthetics during surgical tissue extraction should also be considered. Sevoflurane, an anesthetic, can reduce NMDA receptor activity (Brosnan & Thiesen, 2012) which has also been shown to trigger spine volume shrinkage in cultures (Alvarez et al., 2007; Nagerl et al., 2004; Stein et al., 2015; Wiegert & Oertner, 2013).

While this study focused on biocytin-filled reconstructions to analyze dendritic spine morphology, future studies could benefit from using GFP-labeled cells, as demonstrated by (Schwarz et al., 2019). GFP labeling could potentially increase throughput and the number of morphological analyses without the laborious filling and staining of individual cells, thereby enabling more detailed investigations into spine dynamics and their implications for human brain function and pathology. However, as shown in this study, spines seem to undergo plastic changes in human organotypic slice cultures (Fig. 6). While this may initially seem like a limitation of this technique, it allows for the direct testing of dendritic spine plasticity in response to experimental manipulation. Additionally, taking advantage of rapid developments in cell-specific viral tools, such as new AAV capsids, promoters, and enhancers (Mich et al., 2021), will enable the investigation of specific cells populations in large numbers.

In conclusion, this study demonstrates the value of using human brain tissue to explore dendritic spine morphology, emphasizing the need for direct human studies to understand the unique aspects of the human brain. Our deep learning approach offers promising advancements for 3D analysis, though further technological improvements and data diversity are crucial. This integration of advanced computational tools with human tissue studies will help uncover sex-specific and condition-specific pathomechanisms in neurological disorders, potentially leading to more effective, tailored therapeutic strategies.

**The authors declare no competing financial interests.**

## Acknowledgement

This work was supported by the “Confocal Microscopy Facility”, a core facility of the Interdisciplinary Center for Clinical Research (IZKF) Aachen within the Faculty of Medicine at RWTH Aachen University. We also thank the *Africa Microscopy Initiative Imaging Centre* (RRID: SCR_025881) and Dr. Michael A. Reiche for their support and contribution to this work.

## Notes

### Competing Interest Statement

The authors have declared no competing interest.

## References

Alvarez, V. A., Ridenour, D. A., & Sabatini, B. L. (2007). Distinct structural and ionotropic roles of NMDA receptors in controlling spine and synapse stability. J Neurosci, 27(28), 7365–7376. 10.1523/JNEUROSCI.0956-07.2007

Alvarez, V. A., & Sabatini, B. L. (2007). Anatomical and physiological plasticity of dendritic spines. Annu Rev Neurosci, 30, 79–97. 10.1146/annurev.neuro.30.051606.094222

Anton-Sanchez, L., Larranaga, P., Benavides-Piccione, R., Fernaud-Espinosa, I., DeFelipe, J., & Bielza, C. (2017). Three-dimensional spatial modeling of spines along dendritic networks in human cortical pyramidal neurons. PLoS One, 12(6), e0180400. 10.1371/journal.pone.0180400

Arellano, J. I., Benavides-Piccione, R., Defelipe, J., & Yuste, R. (2007). Ultrastructure of dendritic spines: correlation between synaptic and spine morphologies. Front Neurosci, 1(1), 131–143. 10.3389/neuro.01.1.1.010.2007

Argunsah, A. O., Erdil, E., Ghani, M. U., Ramiro-Cortes, Y., Hobbiss, A. F., Karayannis, T., Cetin, M., Israely, I., & Unay, D. (2022). An interactive time series image analysis software for dendritic spines. Sci Rep, 12(1), 12405. 10.1038/s41598-022-16137-y

Bak, A., Koch, H., van Loo, K. M. J., Schmied, K., Gittel, B., Weber, Y., Ort, J., Schwarz, N., Tauber, S. C., Wuttke, T. V., & Delev, D. (2024). Human organotypic brain slice cultures: a detailed and improved protocol for preparation and long-term maintenance. J Neurosci Methods, 404, 110055. 10.1016/j.jneumeth.2023.110055

Basu, S., Saha, P. K., Roszkowska, M., Magnowska, M., Baczynska, E., Das, N., Plewczynski, D., & Wlodarczyk, J. (2018). Quantitative 3-D morphometric analysis of individual dendritic spines. Sci Rep, 8(1), 3545. 10.1038/s41598-018-21753-8

Beaulieu-Laroche, L., Toloza, E. H. S., van der Goes, M. S., Lafourcade, M., Barnagian, D., Williams, Z. M., Eskandar, E. N., Frosch, M. P., Cash, S. S., & Harnett, M. T. (2018). Enhanced Dendritic Compartmentalization in Human Cortical Neurons. Cell, 175(3), 643–651 e614. 10.1016/j.cell.2018.08.045

Benavides-Piccione, R., Ballesteros-Yanez, I., DeFelipe, J., & Yuste, R. (2002). Cortical area and species differences in dendritic spine morphology. J Neurocytol, 31(3-5), 337–346. 10.1023/a:1024134312173

Benavides-Piccione, R., Fernaud-Espinosa, I., Robles, V., Yuste, R., & DeFelipe, J. (2013). Age-based comparison of human dendritic spine structure using complete three-dimensional reconstructions. Cereb Cortex, 23(8), 1798–1810. 10.1093/cercor/bhs154

Berg, J., Sorensen, S. A., Ting, J. T., Miller, J. A., Chartrand, T., Buchin, A., Bakken, T. E., Budzillo, A., Dee, N., Ding, S. L., Gouwens, N. W., Hodge, R. D., Kalmbach, B., Lee, C., Lee, B. R., Alfiler, L., Baker, K., Barkan, E., Beller, A., . . . Lein, E. S. (2021). Human neocortical expansion involves glutamatergic neuron diversification. Nature, 598(7879), 151–158. 10.1038/s41586-021-03813-8

Berry, K. P., & Nedivi, E. (2017). Spine Dynamics: Are They All the Same? Neuron, 96(1), 43–55. 10.1016/j.neuron.2017.08.008

Boros, B. D., Greathouse, K. M., Gearing, M., & Herskowitz, J. H. (2019). Dendritic spine remodeling accompanies Alzheimer’s disease pathology and genetic susceptibility in cognitively normal aging. Neurobiol Aging, 73, 92–103. 10.1016/j.neurobiolaging.2018.09.003

Bourne, J., & Harris, K. M. (2007). Do thin spines learn to be mushroom spines that remember? Curr Opin Neurobiol, 17(3), 381–386. 10.1016/j.conb.2007.04.009

Bourne, J. N., & Harris, K. M. (2008). Balancing structure and function at hippocampal dendritic spines. Annu Rev Neurosci, 31, 47–67. 10.1146/annurev.neuro.31.060407.125646

Brandt, N., Loffler, T., Fester, L., & Rune, G. M. (2020). Sex-specific features of spine densities in the hippocampus. Sci Rep, 10(1), 11405. 10.1038/s41598-020-68371-x

Brosnan, R. J., & Thiesen, R. (2012). Increased NMDA receptor inhibition at an increased Sevoflurane MAC. BMC Anesthesiol, 12, 9. 10.1186/1471-2253-12-9

Chen, J. R., Wang, T. J., Lim, S. H., Wang, Y. J., & Tseng, G. F. (2013). Testosterone modulation of dendritic spines of somatosensory cortical pyramidal neurons. Brain Struct Funct, 218(6), 1407–1417. 10.1007/s00429-012-0465-7

Chen, M., Qi, J., Poo, M., & Yang, Y. (2022). Stability and dynamics of dendritic spines in macaque prefrontal cortex. Natl Sci Rev, 9(9), nwac125. 10.1093/nsr/nwac125

Chen, X., Leischner, U., Rochefort, N. L., Nelken, I., & Konnerth, A. (2011). Functional mapping of single spines in cortical neurons in vivo. Nature, 475(7357), 501–505. 10.1038/nature10193

Cirelli, C., & Tononi, G. (2020). Effects of sleep and waking on the synaptic ultrastructure. Philos Trans R Soc Lond B Biol Sci, 375(1799), 20190235. 10.1098/rstb.2019.0235

Defelipe, J. (2011). The evolution of the brain, the human nature of cortical circuits, and intellectual creativity. Front Neuroanat, 5, 29. 10.3389/fnana.2011.00029

Delevich, K., Okada, N. J., Rahane, A., Zhang, Z., Hall, C. D., & Wilbrecht, L. (2020). Sex and Pubertal Status Influence Dendritic Spine Density on Frontal Corticostriatal Projection Neurons in Mice. Cereb Cortex, 30(6), 3543–3557. 10.1093/cercor/bhz325

Dhir, N., Medhi, B., Prakash, A., Goyal, M. K., Modi, M., & Mohindra, S. (2020). Pre-clinical to Clinical Translational Failures and Current Status of Clinical Trials in Stroke Therapy: A Brief Review. Curr Neuropharmacol, 18(7), 596–612. 10.2174/1570159X18666200114160844

Duan, H., Wearne, S. L., Rocher, A. B., Macedo, A., Morrison, J. H., & Hof, P. R. (2003). Age-related dendritic and spine changes in corticocortically projecting neurons in macaque monkeys. Cereb Cortex, 13(9), 950–961. 10.1093/cercor/13.9.950

Eyal, G., Verhoog, M. B., Testa-Silva, G., Deitcher, Y., Lodder, J. C., Benavides-Piccione, R., Morales, J., DeFelipe, J., de Kock, C. P., Mansvelder, H. D., & Segev, I. (2016). Unique membrane properties and enhanced signal processing in human neocortical neurons. Elife, 5. 10.7554/eLife.16553

Fiala, J. C., Spacek, J., & Harris, K. M. (2002). Dendritic spine pathology: cause or consequence of neurological disorders? Brain Res Brain Res Rev, 39(1), 29–54. 10.1016/s0165-0173(02)00158-3

Filipek, P. A., Richelme, C., Kennedy, D. N., & Caviness, V. S., Jr. (1994). The young adult human brain: an MRI-based morphometric analysis. Cereb Cortex, 4(4), 344–360. 10.1093/cercor/4.4.344

Franke, C. L., Palm, R., Dalby, M., Schoonderwaldt, H. C., Hantson, L., Eriksson, B., Lang-Jenssen, L., & Smakman, J. (1996). Flunarizine in stroke treatment (FIST): a double-blind, placebo-controlled trial in Scandinavia and the Netherlands. Acta Neurol Scand, 93(1), 56–60. 10.1111/j.1600-0404.1996.tb00171.x

Garelick, T., & Swann, J. (2014). Testosterone regulates the density of dendritic spines in the male preoptic area. Horm Behav, 65(3), 249–253. 10.1016/j.yhbeh.2014.01.008

Ghani, M. U., Mesadi, F., Kanik, S. D., Argunsah, A. O., Hobbiss, A. F., Israely, I., Unay, D., Tasdizen, T., & Cetin, M. (2017). Dendritic spine classification using shape and appearance features based on two-photon microscopy. J Neurosci Methods, 279, 13–21. 10.1016/j.jneumeth.2016.12.006

Gidon, A., Zolnik, T. A., Fidzinski, P., Bolduan, F., Papoutsi, A., Poirazi, P., Holtkamp, M., Vida, I., & Larkum, M. E. (2020). Dendritic action potentials and computation in human layer 2/3 cortical neurons. Science, 367(6473), 83–87. 10.1126/science.aax6239

Gilles, J. F., Mailly, P., Ferreira, T., Boudier, T., & Heck, N. (2024). Spot Spine, a freely available ImageJ plugin for 3D detection and morphological analysis of dendritic spines. F1000Res, 13, 176. 10.12688/f1000research.146327.1

Goldstein, J. M., Seidman, L. J., Horton, N. J., Makris, N., Kennedy, D. N., Caviness, V. S., Jr., Faraone, S. V., & Tsuang, M. T. (2001). Normal sexual dimorphism of the adult human brain assessed by in vivo magnetic resonance imaging. Cereb Cortex, 11(6), 490–497. 10.1093/cercor/11.6.490

Gooch, H. M., Bluett, T., Perumal, M. B., Vo, H. D., Fletcher, L. N., Papacostas, J., Jeffree, R. L., Wood, M., Colditz, M. J., McMillen, J., Tsahtsarlis, T., Amato, D., Campbell, R., Gillinder, L., & Williams, S. R. (2022). High-fidelity dendritic sodium spike generation in human layer 2/3 neocortical pyramidal neurons. Cell Rep, 41(3), 111500. 10.1016/j.celrep.2022.111500

Guidetti, P., Charles, V., Chen, E. Y., Reddy, P. H., Kordower, J. H., Whetsell, W. O., Jr., Schwarcz, R., & Tagle, D. A. (2001). Early degenerative changes in transgenic mice expressing mutant huntingtin involve dendritic abnormalities but no impairment of mitochondrial energy production. Exp Neurol, 169(2), 340–350. 10.1006/exnr.2000.7626

Harris, K. D., & Shepherd, G. M. (2015). The neocortical circuit: themes and variations. Nat Neurosci, 18(2), 170–181. 10.1038/nn.3917

Harris, K. M., Jensen, F. E., & Tsao, B. (1992). Three-dimensional structure of dendritic spines and synapses in rat hippocampus (CA1) at postnatal day 15 and adult ages: implications for the maturation of synaptic physiology and long-term potentiation. J Neurosci, 12(7), 2685–2705. 10.1523/JNEUROSCI.12-07-02685.1992

Hayashi, Y., & Majewska, A. K. (2005). Dendritic spine geometry: functional implication and regulation. Neuron, 46(4), 529–532. 10.1016/j.neuron.2005.05.006

Hunt, S., Leibner, Y., Mertens, E. J., Barros-Zulaica, N., Kanari, L., Heistek, T. S., Karnani, M. M., Aardse, R., Wilbers, R., Heyer, D. B., Goriounova, N. A., Verhoog, M. B., Testa-Silva, G., Obermayer, J., Versluis, T., Benavides-Piccione, R., de Witt-Hamer, P., Idema, S., Noske, D. P., de Kock, C. P. J. (2023). Strong and reliable synaptic communication between pyramidal neurons in adult human cerebral cortex. Cereb Cortex, 33(6), 2857–2878. 10.1093/cercor/bhac246

Iascone, D. M., Li, Y., Sumbul, U., Doron, M., Chen, H., Andreu, V., Goudy, F., Blockus, H., Abbott, L. F., Segev, I., Peng, H., & Polleux, F. (2020). Whole-Neuron Synaptic Mapping Reveals Spatially Precise Excitatory/Inhibitory Balance Limiting Dendritic and Somatic Spiking. Neuron, 106(4), 566–578 e568. 10.1016/j.neuron.2020.02.015

Jabra, S., Rietsche, M., Muellerleile, J., O’Leary, A., Slattery, D. A., Deller, T., & Fellenz, M. (2024). Sex-and cycle-dependent changes in spine density and size in hippocampal CA2 neurons. Sci Rep, 14(1), 12252. 10.1038/s41598-024-62951-x

Jacobs, B., Driscoll, L., & Schall, M. (1997). Life-span dendritic and spine changes in areas 10 and 18 of human cortex: a quantitative Golgi study. J Comp Neurol, 386(4), 661–680. https://www.ncbi.nlm.nih.gov/pubmed/9378859

Kabaso, D., Coskren, P. J., Henry, B. I., Hof, P. R., & Wearne, S. L. (2009). The electrotonic structure of pyramidal neurons contributing to prefrontal cortical circuits in macaque monkeys is significantly altered in aging. Cereb Cortex, 19(10), 2248–2268. 10.1093/cercor/bhn242

Konopaske, G. T., Lange, N., Coyle, J. T., & Benes, F. M. (2014). Prefrontal cortical dendritic spine pathology in schizophrenia and bipolar disorder. JAMA Psychiatry, 71(12), 1323–1331. 10.1001/jamapsychiatry.2014.1582

Koyama, Y., Hattori, T., Nishida, T., Hori, O., & Tohyama, M. (2015). Alterations in dendrite and spine morphology of cortical pyramidal neurons in DISC1-binding zinc finger protein (DBZ) knockout mice. Front Neuroanat, 9, 52. 10.3389/fnana.2015.00052

Lee, A. T., Chang, E. F., Paredes, M. F., & Nowakowski, T. J. (2024). Large-scale neurophysiology and single-cell profiling in human neuroscience. Nature, 630(8017), 587–595. 10.1038/s41586-024-07405-0

Lee, B. R., Dalley, R., Miller, J. A., Chartrand, T., Close, J., Mann, R., Mukora, A., Ng, L., Alfiler, L., Baker, K., Bertagnolli, D., Brouner, K., Casper, T., Csajbok, E., Donadio, N., Driessens, S. L. W., Egdorf, T., Enstrom, R., Galakhova, A. A., . . . Ting, J. T. (2023). Signature morphoelectric properties of diverse GABAergic interneurons in the human neocortex. Science, 382(6667), eadf6484. 10.1126/science.adf6484

Lenz, M., Kruse, P., Eichler, A., Straehle, J., Hemeling, H., Stohr, P., Beck, J., & Vlachos, A. (2024). Clinical parameters affect the structure and function of superficial pyramidal neurons in the adult human neocortex. Brain Commun, 6(5), fcae351. 10.1093/braincomms/fcae351

Levet, F., Tonnesen, J., Nagerl, U. V., & Sibarita, J. B. (2020). SpineJ: A software tool for quantitative analysis of nanoscale spine morphology. Methods, 174, 49–55. 10.1016/j.ymeth.2020.01.020

Loewenstein, Y., Yanover, U., & Rumpel, S. (2015). Predicting the Dynamics of Network Connectivity in the Neocortex. J Neurosci, 35(36), 12535–12544. 10.1523/JNEUROSCI.2917-14.2015

Macdonald, R. L., Higashida, R. T., Keller, E., Mayer, S. A., Molyneux, A., Raabe, A., Vajkoczy, P., Wanke, I., Bach, D., Frey, A., Marr, A., Roux, S., & Kassell, N. (2011). Clazosentan, an endothelin receptor antagonist, in patients with aneurysmal subarachnoid haemorrhage undergoing surgical clipping: a randomised, double-blind, placebo-controlled phase 3 trial (CONSCIOUS-2). Lancet Neurol, 10(7), 618–625. 10.1016/S1474-4422(11)70108-9

Malinow, R., & Malenka, R. C. (2002). AMPA receptor trafficking and synaptic plasticity. Annu Rev Neurosci, 25, 103–126. 10.1146/annurev.neuro.25.112701.142758

Matsuzaki, M., Honkura, N., Ellis-Davies, G. C., & Kasai, H. (2004). Structural basis of long-term potentiation in single dendritic spines. Nature, 429(6993), 761–766. 10.1038/nature02617

Mich, J. K., Graybuck, L. T., Hess, E. E., Mahoney, J. T., Kojima, Y., Ding, Y., Somasundaram, S., Miller, J. A., Kalmbach, B. E., Radaelli, C., Gore, B. B., Weed, N., Omstead, V., Bishaw, Y., Shapovalova, N. V., Martinez, R. A., Fong, O., Yao, S., Mortrud, M., . . . Levi, B. P. (2021). Functional enhancer elements drive subclass-selective expression from mouse to primate neocortex. Cell Rep, 34(13), 108754. 10.1016/j.celrep.2021.108754

Mohan, H., Verhoog, M. B., Doreswamy, K. K., Eyal, G., Aardse, R., Lodder, B. N., Goriounova, N. A., Asamoah, B., AB, B. B., Groot, C., van der Sluis, S., Testa-Silva, G., Obermayer, J., Boudewijns, Z. S., Narayanan, R. T., Baayen, J. C., Segev, I., Mansvelder, H. D., & de Kock, C. P. (2015). Dendritic and Axonal Architecture of Individual Pyramidal Neurons across Layers of Adult Human Neocortex. Cereb Cortex, 25(12), 4839–4853. 10.1093/cercor/bhv188

Molnar, G., Rozsa, M., Baka, J., Holderith, N., Barzo, P., Nusser, Z., & Tamas, G. (2016). Human pyramidal to interneuron synapses are mediated by multi-vesicular release and multiple docked vesicles. Elife, 5. 10.7554/eLife.18167

Motley, S. E., Grossman, Y. S., Janssen, W. G. M., Baxter, M. G., Rapp, P. R., Dumitriu, D., & Morrison, J. H. (2018). Selective Loss of Thin Spines in Area 7a of the Primate Intraparietal Sulcus Predicts Age-Related Working Memory Impairment. J Neurosci, 38(49), 10467–10478. 10.1523/JNEUROSCI.1234-18.2018

Musall, S., Sun, X. R., Mohan, H., An, X., Gluf, S., Li, S. J., Drewes, R., Cravo, E., Lenzi, I., Yin, C., Kampa, B. M., & Churchland, A. K. (2023). Pyramidal cell types drive functionally distinct cortical activity patterns during decision-making. Nat Neurosci, 26(3), 495–505. 10.1038/s41593-022-01245-9

Nagerl, U. V., Eberhorn, N., Cambridge, S. B., & Bonhoeffer, T. (2004). Bidirectional activity-dependent morphological plasticity in hippocampal neurons. Neuron, 44(5), 759–767. 10.1016/j.neuron.2004.11.016

Nimchinsky, E. A., Sabatini, B. L., & Svoboda, K. (2002). Structure and function of dendritic spines. Annu Rev Physiol, 64, 313–353. 10.1146/annurev.physiol.64.081501.160008

Nusser, Z., Lujan, R., Laube, G., Roberts, J. D., Molnar, E., & Somogyi, P. (1998). Cell type and pathway dependence of synaptic AMPA receptor number and variability in the hippocampus. Neuron, 21(3), 545–559. 10.1016/s0896-6273(00)80565-6

Page, T. L., Einstein, M., Duan, H., He, Y., Flores, T., Rolshud, D., Erwin, J. M., Wearne, S. L., Morrison, J. H., & Hof, P. R. (2002). Morphological alterations in neurons forming corticocortical projections in the neocortex of aged Patas monkeys. Neurosci Lett, 317(1), 37–41. 10.1016/s0304-3940(01)02428-4

Papa, M., Bundman, M. C., Greenberger, V., & Segal, M. (1995). Morphological analysis of dendritic spine development in primary cultures of hippocampal neurons. J Neurosci, *15*(1 Pt 1), 1-11. 10.1523/JNEUROSCI.15-01-00001.1995

Passe, T. J., Rajagopalan, P., Tupler, L. A., Byrum, C. E., MacFall, J. R., & Krishnan, K. R. (1997). Age and sex effects on brain morphology. Prog Neuropsychopharmacol Biol Psychiatry, 21(8), 1231–1237. 10.1016/s0278-5846(97)00160-7

Peng, Y., Bjelde, A., Aceituno, P. V., Mittermaier, F. X., Planert, H., Grosser, S., Onken, J., Faust, K., Kalbhenn, T., Simon, M., Radbruch, H., Fidzinski, P., Schmitz, D., Alle, H., Holtkamp, M., Vida, I., Grewe, B. F., & Geiger, J. R. P. (2024). Directed and acyclic synaptic connectivity in the human layer 2-3 cortical microcircuit. Science, 384(6693), 338–343. 10.1126/science.adg8828

Penzes, P., Cahill, M. E., Jones, K. A., VanLeeuwen, J. E., & Woolfrey, K. M. (2011). Dendritic spine pathology in neuropsychiatric disorders. Nat Neurosci, 14(3), 285–293. 10.1038/nn.2741

Petanjek, Z., Judas, M., Simic, G., Rasin, M. R., Uylings, H. B., Rakic, P., & Kostovic, I. (2011). Extraordinary neoteny of synaptic spines in the human prefrontal cortex. Proc Natl Acad Sci U S A, 108(32), 13281–13286. 10.1073/pnas.1105108108

Peters, A., Sethares, C., & Moss, M. B. (1998). The effects of aging on layer 1 in area 46 of prefrontal cortex in the rhesus monkey. Cereb Cortex, 8(8), 671–684. 10.1093/cercor/8.8.671

Rada, L., Kilic, B., Erdil, E., Ramiro-Cortes, Y., Israely, I., Unay, D., Cetin, M., & Argunsah, A. O. (2018). Tracking-assisted Detection of Dendritic Spines in Time-Lapse Microscopic Images. Neuroscience, 394, 189–205. 10.1016/j.neuroscience.2018.10.022

Reberger, R., Dall’Oglio, A., Jung, C. R., & Rasia-Filho, A. A. (2018). Structure and diversity of human dendritic spines evidenced by a new three-dimensional reconstruction procedure for Golgi staining and light microscopy. J Neurosci Methods, 293, 27–36. 10.1016/j.jneumeth.2017.09.001

Roberts, R. C., Conley, R., Kung, L., Peretti, F. J., & Chute, D. J. (1996). Reduced striatal spine size in schizophrenia: a postmortem ultrastructural study. Neuroreport, 7(6), 1214–1218. 10.1097/00001756-199604260-00024

Rochefort, N. L., & Konnerth, A. (2012). Dendritic spines: from structure to in vivo function. EMBO Rep, 13(8), 699–708. 10.1038/embor.2012.102

Salminen, L. E., Tubi, M. A., Bright, J., Thomopoulos, S. I., Wieand, A., & Thompson, P. M. (2022). Sex is a defining feature of neuroimaging phenotypes in major brain disorders. Hum Brain Mapp, 43(1), 500–542. 10.1002/hbm.25438

Schwarz, N., Uysal, B., Welzer, M., Bahr, J. C., Layer, N., Loffler, H., Stanaitis, K., Pa, H., Weber, Y. G., Hedrich, U. B., Honegger, J. B., Skodras, A., Becker, A. J., Wuttke, T. V., & Koch, H. (2019). Long-term adult human brain slice cultures as a model system to study human CNS circuitry and disease. Elife, 8. 10.7554/eLife.48417

Segal, M. (2017). Dendritic spines: Morphological building blocks of memory. Neurobiol Learn Mem, 138, 3–9. 10.1016/j.nlm.2016.06.007

Seyhan, A. A. (2019). Lost in translation: the valley of death across preclinical and clinical divide – identification of problems and overcoming obstacles. Translational Medicine Communications, 4(1), 18. 10.1186/s41231-019-0050-7

Shi, P., Huang, Y., & Hong, J. (2014). Automated three-dimensional reconstruction and morphological analysis of dendritic spines based on semi-supervised learning. Biomed Opt Express, 5(5), 1541–1553. 10.1364/BOE.5.001541

Shors, T. J., Chua, C., & Falduto, J. (2001). Sex differences and opposite effects of stress on dendritic spine density in the male versus female hippocampus. J Neurosci, 21(16), 6292–6297. 10.1523/JNEUROSCI.21-16-06292.2001

Smirnov, M. S., Garrett, T. R., & Yasuda, R. (2018). An open-source tool for analysis and automatic identification of dendritic spines using machine learning. PLoS One, 13(7), e0199589. 10.1371/journal.pone.0199589

Stein, I. S., Gray, J. A., & Zito, K. (2015). Non-Ionotropic NMDA Receptor Signaling Drives Activity-Induced Dendritic Spine Shrinkage. J Neurosci, 35(35), 12303–12308. 10.1523/JNEUROSCI.4289-14.2015

Stuart, G. J., & Sakmann, B. (1994). Active propagation of somatic action potentials into neocortical pyramidal cell dendrites. Nature, 367(6458), 69–72. 10.1038/367069a0

Swanger, S. A., Yao, X., Gross, C., & Bassell, G. J. (2011). Automated 4D analysis of dendritic spine morphology: applications to stimulus-induced spine remodeling and pharmacological rescue in a disease model. Mol Brain, 4, 38. 10.1186/1756-6606-4-38

Tazerart, S., Mitchell, D. E., Miranda-Rottmann, S., & Araya, R. (2020). A spike-timing-dependent plasticity rule for dendritic spines. Nat Commun, 11(1), 4276. 10.1038/s41467-020-17861-7

Ting, J. T., Kalmbach, B., Chong, P., de Frates, R., Keene, C. D., Gwinn, R. P., Cobbs, C., Ko, A. L., Ojemann, J. G., Ellenbogen, R. G., Koch, C., & Lein, E. (2018). A robust ex vivo experimental platform for molecular-genetic dissection of adult human neocortical cell types and circuits. Sci Rep, 8(1), 8407. 10.1038/s41598-018-26803-9

Tonnesen, J., Katona, G., Rozsa, B., & Nagerl, U. V. (2014). Spine neck plasticity regulates compartmentalization of synapses. Nat Neurosci, 17(5), 678–685. 10.1038/nn.3682

Vardalaki, D., Chung, K., & Harnett, M. T. (2022). Filopodia are a structural substrate for silent synapses in adult neocortex. Nature, 612(7939), 323–327. 10.1038/s41586-022-05483-6

Verhoog, M. B., Goriounova, N. A., Obermayer, J., Stroeder, J., Hjorth, J. J., Testa-Silva, G., Baayen, J. C., de Kock, C. P., Meredith, R. M., & Mansvelder, H. D. (2013). Mechanisms underlying the rules for associative plasticity at adult human neocortical synapses. J Neurosci, 33(43), 17197–17208. 10.1523/JNEUROSCI.3158-13.2013

Vidaurre-Gallart, I., Fernaud-Espinosa, I., Cosmin-Toader, N., Talavera-Martinez, L., Martin-Abadal, M., Benavides-Piccione, R., Gonzalez-Cid, Y., Pastor, L., DeFelipe, J., & Garcia-Lorenzo, M. (2022). A Deep Learning-Based Workflow for Dendritic Spine Segmentation. Front Neuroanat, 16, 817903. 10.3389/fnana.2022.817903

Vogel, F. W., Alipek, S., Eppler, J. B., Osuna-Vargas, P., Triesch, J., Bissen, D., Acker-Palmer, A., Rumpel, S., & Kaschube, M. (2023). Utilizing 2D-region-based CNNs for automatic dendritic spine detection in 3D live cell imaging. Sci Rep, 13(1), 20497. 10.1038/s41598-023-47070-3

Wahlgren, N. G., Ranasinha, K. W., Rosolacci, T., Franke, C. L., van Erven, P. M., Ashwood, T., & Claesson, L. (1999). Clomethiazole acute stroke study (CLASS): results of a randomized, controlled trial of clomethiazole versus placebo in 1360 acute stroke patients. Stroke, 30(1), 21–28. 10.1161/01.str.30.1.21

Wefelmeyer, W., Puhl, C. J., & Burrone, J. (2016). Homeostatic Plasticity of Subcellular Neuronal Structures: From Inputs to Outputs. Trends Neurosci, 39(10), 656–667. 10.1016/j.tins.2016.08.004

Wiegert, J. S., & Oertner, T. G. (2013). Long-term depression triggers the selective elimination of weakly integrated synapses. Proc Natl Acad Sci U S A, 110(47), E4510–4519. 10.1073/pnas.1315926110

Woolley, C. S., Gould, E., Frankfurt, M., & McEwen, B. S. (1990). Naturally occurring fluctuation in dendritic spine density on adult hippocampal pyramidal neurons. J Neurosci, 10(12), 4035–4039. 10.1523/JNEUROSCI.10-12-04035.1990

Xiao, X., Djurisic, M., Hoogi, A., Sapp, R. W., Shatz, C. J., & Rubin, D. L. (2018). Automated dendritic spine detection using convolutional neural networks on maximum intensity projected microscopic volumes. J Neurosci Methods, 309, 25–34. 10.1016/j.jneumeth.2018.08.019

Yang, D., Qi, G., Ort, J., Witzig, V., Bak, A., Delev, D., Koch, H., & Feldmeyer, D. (2024). Modulation of large rhythmic depolarizations in human large basket cells by norepinephrine and acetylcholine. Commun Biol, 7(1), 885. 10.1038/s42003-024-06546-2

Young, M. E., Ohm, D. T., Dumitriu, D., Rapp, P. R., & Morrison, J. H. (2014). Differential effects of aging on dendritic spines in visual cortex and prefrontal cortex of the rhesus monkey. Neuroscience, 274, 33–43. 10.1016/j.neuroscience.2014.05.008

Yuste, R., & Bonhoeffer, T. (2004). Genesis of dendritic spines: insights from ultrastructural and imaging studies. Nat Rev Neurosci, 5(1), 24–34. 10.1038/nrn1300

Zhou, Q., Homma, K. J., & Poo, M. M. (2004). Shrinkage of dendritic spines associated with long-term depression of hippocampal synapses. Neuron, 44(5), 749–757. 10.1016/j.neuron.2004.11.011

Zhou, Y., Lai, C. S. W., Bai, Y., Li, W., Zhao, R., Yang, G., Frank, M. G., & Gan, W. B. (2020). REM sleep promotes experience-dependent dendritic spine elimination in the mouse cortex. Nat Commun, 11(1), 4819. 10.1038/s41467-020-18592-5

